# Genome-wide ancestry and introgression in a Zambian baboon hybrid zone

**DOI:** 10.1101/578781

**Authors:** Kenneth L. Chiou, Christina M. Bergey, Andrew S. Burrell, Todd R. Disotell, Jeffrey Rogers, Clifford J. Jolly, Jane E. Phillips-Conroy

## Abstract

Hybridization in nature offers unique insights into the process of natural selection in incipient species and their hybrids. In order to evaluate the patterns and targets of selection, we examine a recently discovered baboon hybrid zone in the Kafue River valley of Zambia, where Kinda baboons (*Papio kindae*) and gray-footed chacma baboons (*P. ursinus griseipes*) coexist with hybridization. We genotyped baboons at 14,962 variable genome-wide autosomal markers using double-digest RADseq. We compare ancestry patterns from this genome-wide dataset to previously reported ancestry from mitochondrial-DNA and Y-chromosome sources. We also fit a Bayesian genomic cline model to scan for genes with extreme patterns of introgression. We show that the Kinda baboon Y chromosome has penetrated the species boundary to a greater extent than either mitochondrial DNA or the autosomal chromosomes. We also find evidence for overall restricted introgression in the JAK/STAT signaling pathway. Echoing results in other species including humans, we find evidence for enhanced and/or directional introgression of immune-related genes or pathways including the toll-like receptor pathway, the blood coagulation pathway, and the *LY96* gene. Finally we show enhanced introgression and excess chacma baboon ancestry in the sperm tail gene *ODF2*. Together, our results elucidate the dynamics of introgressive hybridization in a primate system while highlighting genes and pathways under selection.

## Introduction

Among many organisms including primates, divergence with reticulate evolution is extremely common to the extent that in many lineages, hybridization during divergence may be considered a rule rather than an exception (Arnold & Meyer 2006; Abbott *et al.* 2013). Hybridization in the wild offers opportune natural systems for evaluating the actions of selection on species-specific traits (Barton & Hewitt 1989; Harrison 1990) including aspects of physiology, morphology, and behavior. At the DNA level, natural selection impacts the patterns by which genetic regions are exchanged between hybridizing lineages, with the direction and rate of introgression influenced by their selective advantage or disadvantage in differentiated genomic backgrounds (Gompert & Buerkle 2011). Regions containing variants that reduce the fitness of hybrids, for example, experience limited penetration of species boundaries while regions containing variants that increase the fitness of hybrids experience more rapid penetration of species boundaries. The latter process characterizes adaptive introgression (Hedrick 2013).

Our present study centers on a hybrid zone between Kinda (*Papio kindae*) and gray-footed chacma baboons (*P. ursinus griseipes*) in the Kafue River valley in central Zambia (Figure S1). Here, the distributions of the two species adjoin in and around the area of Kafue National Park, with Kinda and chacma baboons occupying respectively the northern and southern areas of the region. After previous speculation regarding potential overlap and interbreeding in the region (Ansell 1960, 1978; Jolly 1993; Burrell 2009), Jolly *et al.* (2011) reported that the two species did in fact hybridize, detecting individuals in multiple localities with mixed or intermediate phenotypes as well as individuals and social groups with contrasting ancestry inferred from mitochondrial DNA and the Y chromosome. The Kafue National Park hybrid zone adds to extensive documentation of past and present hybridization in baboons (Jolly 1993; Zinner *et al.* 2011; Rogers *et al.* 2019), including ongoing hybridization around Awash National Park in Ethiopia (Phillips-Conroy & Jolly 1986; Bergman & Beehner 2004), Amboseli National Park in Kenya (Alberts & Altmann 2001; Tung *et al.* 2008; Charpentier *et al.* 2012), and Gorongosa National Park in Mozambique (Martinez *et al.* 2019).

Hybridization in the Kafue River valley is striking given unusually pronounced body-size differences between the two species (Jolly *et al.* 2011). Gray-footed chacma baboons are among the largest baboon species, second only to southern Cape chacma baboons (*P. ursinus ursinus*). Kinda baboons, on the other hand, are the smallest baboon species, with adult males and females weighing approximately 53% and 74% of the body mass of their respective gray-footed chacma baboon counterparts. In a typical male gray-footed chacma × female Kinda baboon mating, the female partner would weigh roughly 35% of the mass of the male partner, while in a typical female gray-footed chacma × male Kinda baboon mating, the female partner would weigh 88% of the mass of the male partner (Jolly *et al.* 2011). Each pairing would match an extreme of body-mass sexual dimorphism in extant papionin monkey species (Delson *et al.* 2000) and would potentially be susceptible to both pre- and post-zygotic obstacles to successful reproduction.

Hybridization in the Kafue River valley is also interesting in the context of behavioral differences between species and the interaction of competing behaviors upon contact (Bergman *et al.* 2008; Charpentier *et al.* 2012). Chacma baboons, like all other baboon species except hamadryas and Guinea baboons (Kummer 1968; Fischer *et al.* 2017), live in multi-male/multi-female social groups characterized by strong female bonds and male dispersal (Silk *et al.* 2009, 2010). Mating is to a high degree determined by both male and female rank (Bulger 1993). Kinda baboons are much less known, but appear also to live in multi-male/multi-female social groups with female bonds and male dispersal (A. Weyher, pers. comm.). Curiously, male Kinda baboons frequently initiate and maintain grooming interactions with non-estrous females, a behavior that is rarely seen in any other baboon species (Weyher *et al.* 2014), suggesting divergent sex-specific strategies for reproduction in this species.

Notably, Jolly *et al.* (2011) found that, among hybrid males with discordant mitochondrial-DNA and Y-chromosome ancestry, the Y chromosome was exclusively inherited from Kinda baboons, a surprising finding given the diminutive size of male Kinda baboons, which would likely be physically outmatched by larger male chacma baboons in the context of highly competitive mating (Bulger 1993). This finding instead suggests that male chacma baboons experienced lower reproductive success during initial hybridization. Jolly *et al.* (2011) hypothesized that this may be a result of either, or both, of two factors: reproductive incompatibilities, potentially related to an obstetric mismatch between small Kinda-like mothers and unusually large hybrid fetuses, or a propensity for surreptitious (“sneak”) matings by male Kinda baboons, facilitated by their immature (i.e., small and lanky) and thereby nonthreatening appearance to male chacma competitors.

In the present study, we employ a population-genomic approach to investigate genome-wide patterns of ancestry and selection in the Kafue River valley hybrid zone. We use double-digest RADseq (Peterson *et al.* 2012) to sequence tens of thousands of genomic variants to high coverage, with additional enrichment of DNA from feces (Chiou & Bergey 2018) to incorporate noninvasive genomic information from unhabituated wild baboons. We use the resulting genomic dataset to characterize patterns of genomic ancestry in the hybrid zone and to identify candidate genes subject to extreme patterns of introgression.

## Methods

### Samples

Biological samples from wild Zambian baboons were collected between 2006 and 2015 from the Kafue River valley contact zone in Zambia (Jolly *et al.* 2011) (Table 1). Blood and fecal samples were collected and stored following procedures described elsewhere (Jolly *et al.* 2011; Chiou & Bergey 2018; Chiou *et al.* 2019). All procedures were conducted following laws and regulations in Zambia and with approval from the Animal Studies Committees at Washington University, New York University, and Baylor College of Medicine. This research follows the International Primatological Society’s Code of Best Practices for Field Primatology.

**Table 1:** Samples included in this analysis. Only samples that passed filters are listed in this table. An expanded list including sample IDs is provided in Table S1.

| Locality | Latitude | Longitude | Year | Appearance | Tissue type | N |
| --- | --- | --- | --- | --- | --- | --- |
| Chunga | -15.0446° | 25.9994° | 2011 | Kinda | blood | 71 |
| Mwengwa Rapids | -15.3115° | 25.9449° | 2014 | mostly Kinda | feces | 5 |
| Malala Camp | -15.7661° | 25.8626° | 2014 | mixed | feces | 3 |
| Musa Bridge | -15.9081° | 25.7414° | 2007, 2015 | mixed | feces | 4 |
| Top Musa | -15.9013° | 25.8133° | 2014 | mixed | feces | 5 |
| North Nkala Road | -15.9029° | 25.8895° | 2007 | mixed | feces | 6 |
| Ngoma Airstrip | -15.9695° | 25.9376° | 2012 | mixed | blood | 12 |
| Dendro Park | -16.1518° | 26.0599° | 2012 | mostly chacma | blood | 4 |
| Nanzhila Plains | -16.2337° | 25.9612° | 2007 | mostly chacma | feces | 5 |
| Choma | -16.6383° | 27.0308° | 2007 | chacma | feces | 3 |
| Lower Zambezi National Park | -15.9234° | 28.9466° | 2006 | chacma | feces | 5 |

### DNA isolation, library preparation and sequencing

We sequenced DNA from a total of 129 animals, of which 123 remained after filtering for quality as described in the following section (Table S1). For blood samples, which included isolated leukocytes, plasma, or FTA-dried blood spots, we extracted DNA using the QIAamp DNA Blood Mini Kit (Qiagen). For fecal samples, we extracted DNA using the QIAamp DNA Stool Mini Kit (Qiagen).

Because DNA in feces is overwhelmingly of bacterial origin (Perry *et al.* 2010; Snyder-Mackler *et al.* 2016), we enriched host DNA from fecal samples using FecalSeq (Chiou & Bergey 2018), which takes advantage of the high density of CpG-methylation in mammalian genomes relative to bacterial genomes to partition fecal DNA into endogenous and exogenous fractions using methyl-binding domain protein baits. We then used double-digest RADseq (ddRADseq) (Peterson *et al.* 2012) to sequence a representative and reproducible set of orthologous SNPs. Procedures for fecal enrichment and ddRADseq library preparation are described elsewhere (Chiou & Bergey 2018; Chiou *et al.* 2019). We sequenced libraries on the Illumina HiSeq 2500 or Illumina HiSeq 4000 platforms.

Bioinformatic procedures for sample demultiplexing, sequence alignment to the anubis baboon reference genome (Panu2.0) (Rogers *et al.* 2019), and variant identification followed methods described elsewhere (Chiou & Bergey 2018; Chiou *et al.* 2019). After calling and filtering variants, we identified and removed variants in strong linkage disequilibrium with PLINK (Purcell *et al.* 2007; Chang *et al.* 2015), using a sliding window approach with a window length of 50 SNPs, a step size of 5 SNPs, and an *r*^2^ threshold of 0.5 (`--indep-pairwise 50 5 0.5`). The final dataset contained 14,642 SNPs.

### Detecting introgression

In an exploratory analysis, we first performed a classical multidimensional scaling (MDS) analysis to visualize identity-by-state using PLINK (Purcell *et al.* 2007; Chang *et al.* 2015) (Figure S2).

We next performed an unsupervised clustering using ADMIXTURE in order to estimate the ancestry of each individual by maximum likelihood (Alexander *et al.* 2009). We ran the algorithm with 10-fold cross validation to estimate the error rate using values for *K* ranging from 1 to 10. We found that *K* = 2 was the most likely value (Figure S3). We also estimated standard errors for point estimates of the *Q* parameter using moving block bootstrapping, after calibrating the dataset such that 0 corresponded to pure Kinda baboon ancestry and 1 corresponded to pure chacma baboon ancestry. We evaluated concordance between the MDS and ADMIXTURE results by linear regression (Figure S4).

We compared genetic ancestry results for the present study to our previously reported ancestry results derived from matrilineal mitochondrial and patrilineal Y-chromosome markers (Jolly *et al.* 2011). As with the autosomal data, we assigned a hybrid index to all individuals that was either 0, indicating a Kinda baboon haplotype, or 1, indicating a chacma baboon haplotype. Because individuals and sampling locales differed between studies, we calculated mean ancestries for each locale and marker set and interpolated the results across geographic space at 1 km^2^ resolution using a Kriging surface model procedure implemented in the fields package (Nychka *et al.* 2015) in R (R Core Team 2013). We then assigned for each sampling locale (*n* = 40) and marker set a single hybrid index based on the geographic interpolation results. To make up for gaps in sampling, we also randomly drew an equal number of “simulated” sampling locales sampled from the geographic background (latitudinal limits: 14.0°S, 17.2°S; longitudinal limits: 25.0°E, 29.2°E) for a total of 80 locales. We then compared ancestry results between marker sets in a pairwise fashion (i.e., across all three possible pairs of marker sets) using nonparametric Wilcoxon signed rank tests. These comparisons were repeated with and without simulated locales.

### Genomic cline analysis

In order to identify regions with outlier patterns of ancestry and introgression, we employed the Bayesian genomic cline method (Gompert & Buerkle 2011) implemented in bgc (Gompert & Buerkle 2012). Bayesian genomic clines provide a robust statistical framework for identifying extreme patterns of introgression by first modeling neutral patterns in the dataset. The model therefore provides a useful method for identifying loci potentially associated with variation in fitness that is calibrated to the provided dataset.

As bgc requires the *a priori* specification of ancestral populations, we first classified individuals as pure or admixed based on results from the ADMIXTURE model described above. We considered individuals to be of pure ancestry if their ancestry estimates inferred by ADMIXTURE were greater than 0.999. By this criterion, we identified 69 pure Kinda baboons, 20 pure chacma baboons, and 34 individuals of mixed ancestry (Figure S5).

We analyzed the final SNP dataset in 5 runs of 400,000 MCMC iterations each, sampling every 40 iterations after discarding the first 200,000 iterations as burn in. We set Kinda baboons as species 0 and chacma baboons as species 1. To incorporate information about genotypic uncertainty, we ran the analysis using raw allelic read counts rather than genotype calls for all 14,642 SNPs in the dataset (arguments: `-O 0 -x 400000 -n 200000 -t 40 -N 1 -E 0.0001 -q 1 -I 1 -p 1`).

Each run was assessed for convergence visually (Figure S6 and Figure S7) and statistically using the Heidelberger & Welch (1983) convergence diagnostic as implemented in the coda package (Plummer *et al.* 2006) in R (R Core Team 2013). We ran both a stationarity test and a half-width test on posterior sample estimates to ensure adequate convergence. The stationarity test uses the Cramer-von-Mises statistic to test a null hypothesis of stationarity on the posterior sample chain with successively higher fractions removed until either the null hypothesis is accepted, indicating convergence, or 50% of the chain has been discarded, indicating failure. We used an α = 0.05 for this test. The half-width test uses the stationary portion of the posterior chain identified previously to calculate the ratio between one-half the width of the 95% confidence interval and the mean. A ratio lower than ε indicates that the sample length is sufficient to estimate the mean accurately while a ratio higher than ε indicates a sample of insufficient length. We set ε = 0.1 for this analysis. For each chain, we excluded loci that did not converge. We then combined MCMC samples from all chains.

From the combined MCMC sample, we drew posterior estimates for the cline parameters *α*_*i*_, which measures the shift of the cline (positive values: shift to species 1; negative values: shift to species 0), and *β*_*i*_, which measures the gradient of the cline (positive values: steep cline; negative values: wide cline). Because protein-coding genes are the focus of the analysis, we summarized parameter estimates by assigning a single value of *α*_*i*_ and *β*_*i*_ to each protein-coding gene. For each MCMC sample, we assigned values of *α*_*i*_ and *β*_*i*_ to each gene calculated as the mean of point parameter estimates for SNPs falling within the boundaries of the protein-coding gene ± 50 bp. We thus derived posterior MCMC samples of cline parameter estimates for a total of 2,249 genes. Genome coordinates were based on the baboon reference genome (Panu2.0) gene annotations (Rogers *et al.* 2019).

In order to test for extreme patterns of introgression of protein-coding genes, we summarized the posterior chains for genes by calculating mean point estimates as well as 95% equal-tail probability (ETP) credible intervals. Following Gompert & Buerkle (2012), we characterized loci as having patterns of excess ancestry if the 95% ETP intervals for *α*_*i*_ or *β*_*i*_ did not contain 0. Candidate loci can thus be interpreted as having at least a 95% probability of having a nonzero value for a particular statistic. Separately, for the purposes of enrichment analyses described in the following section, we calculated *p*-value analogs for each Bayesian statistic, defined as 1 minus the fraction of posterior point estimates greater than (for testing positive values) or less than 0 (for testing negative values). The genes identified on the basis of our calculated *p* analogs (*p* < 0.025) and 95% ETP intervals are equivalent.

### Functional enrichment analysis

Instead of testing for significant or extreme signatures associated with single genes, enrichment analysis tests for extreme patterns associated overall with well-characterized biological functions or pathways. Enrichment analysis may therefore detect broadly shared but subtle effects of evolution on biologically relevant gene sets, even when few or none of the associated genes are individually identified as extreme.

We conducted enrichment analyses at two levels: biological processes from the Gene Ontology (GO) (Gene Ontology Consortium 2000, 2015) and biological pathways from the PANTHER pathway database (Mi *et al.* 2013b; a). We downloaded GO annotations associated with the baboon reference genome (Panu2.0) using biomaRt (Durinck *et al.* 2009). Because PANTHER annotations do not yet exist for the baboon genome, we derived pathway annotations through homology from the rhesus macaque proteome following methods described elsewhere (Chiou *et al.* 2019).

For our GO enrichment analyses, we tested for directional enrichment of *α*_*i*_ or *β*_*i*_ using a modified Kolmogorov–Smirnov test implemented in topGO (Alexa & Rahnenführer 2018). This procedure increases biological signal by taking into account the underlying hierarchical topology of GO terms (Alexa *et al.* 2006). In order to incorporate information on the variance of posterior point estimates, rather than simply the mean, we ran the test using our calculated *p* analogs describe above. We ran the “weight01” algorithm, after limiting nodes to those in the “biological process” ontology and those associated with at least 10 annotated genes in our dataset. We considered terms with a *p*-value < 0.025 (Bonferroni-adjusted for two one-sided tests) to be enriched for extreme *α* or *β*. Correction for testing across multiple terms was not conducted due to the exploratory nature of these analyses.

For our PANTHER enrichment analysis, we used Wilcoxon rank sum tests to evaluate, for each pathway, whether *p* analogs assigned to each pathway were more extreme than values not assigned to that pathway. We performed for each statistic a pair of one-tailed tests to scan for pathways associated with negative or positive *α*_*i*_ or *β*_*i*_ values. As with the GO analyses, we considered terms with a *p*-value < 0.025 (Bonferroni-adjusted for two one-sided tests) to be enriched for extreme *α* or *β* and did not correct for multiple hypothesis testing. All enrichment analyses were run in R (R Core Team 2013).

## Results

We used double-digest RADseq (Peterson *et al.* 2012) in combination with FecalSeq enrichment of the host genome for an important subset of DNA samples derived from feces (Chiou & Bergey 2018; Chiou *et al.* 2019) to genotype Zambian baboons from the greater Kafue National Park region at thousands of polymorphic autosomal sites. After filtering out loci with insufficient quality and high missingness, as well as loci in high linkage disequilibrium, we derived a final dataset of 123 Kinda, chacma, and hybrid baboons genotyped at 14,962 SNP loci. The mean sequencing depth across these loci was 41.94 reads per individual, but varied across individuals from 0.29 to 233.66, with 90% of individuals sequenced to an average of at least 5.23 reads per locus.

### Hybrid zone structure

We investigated population structure and estimated ancestry components using the unsupervised maximum likelihood population clustering approach implemented in ADMIXTURE (Alexander *et al.* 2009) (Figure 1 and Table S1). We used the *Q* parameters from the model as estimates of a hybrid index ranging from 0 to 1 for each individual, with 0 indicating pure Kinda baboon ancestry and 1 indicating pure chacma baboon ancestry.

**Fig 1:**
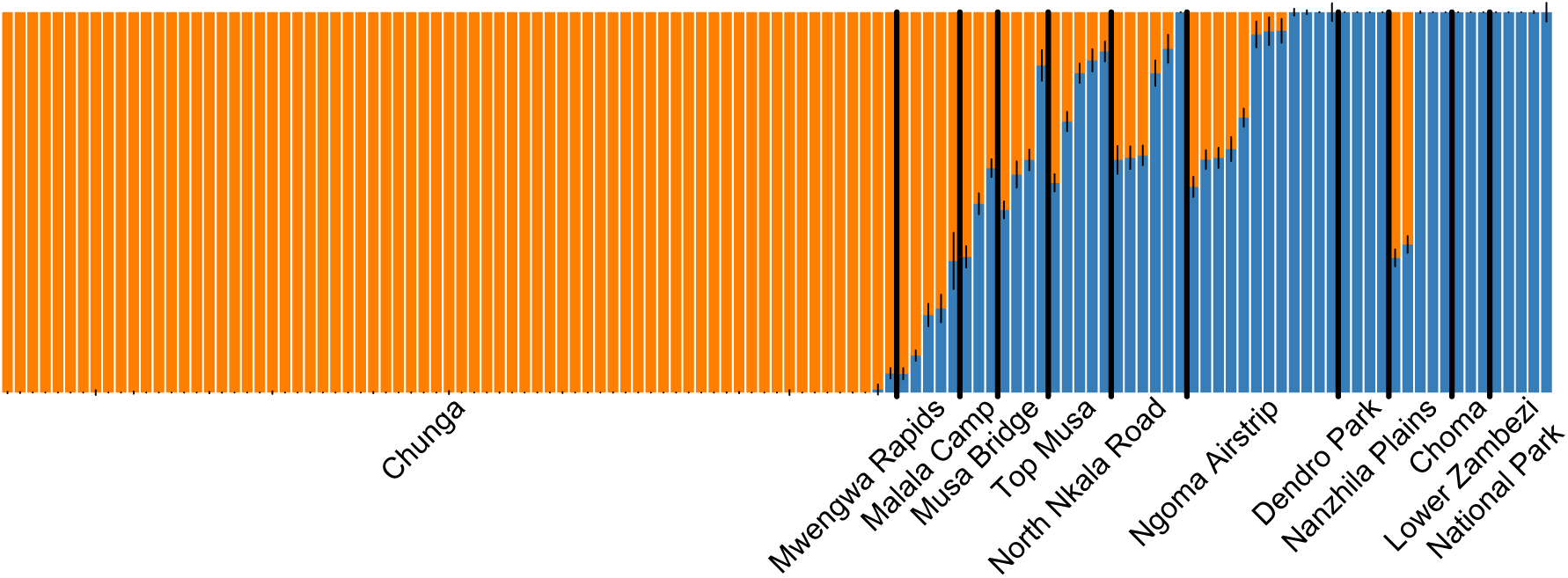
Ancestry estimation results using ADMIXTURE. Individuals are ordered according to the localities listed in Table 1.

The ancestry results when visualized over geographic space revealed a wide cline, spanning at least 100 km on a north-south axis (Figure 2). As expected based on their location and physical appearance, individuals from Chunga, located north of the hybrid zone (Ansell 1978; Jolly *et al.* 2011), were almost exclusively assigned pure Kinda baboon ancestry while individuals from Choma and Lower Zambezi National Park, located well south and east of the Kinda baboon distribution (Ansell 1978), were almost exclusively assigned pure chacma baboon ancestry. These findings corroborate previous results based on microsatellite markers (Burrell 2009).

**Fig 2:**
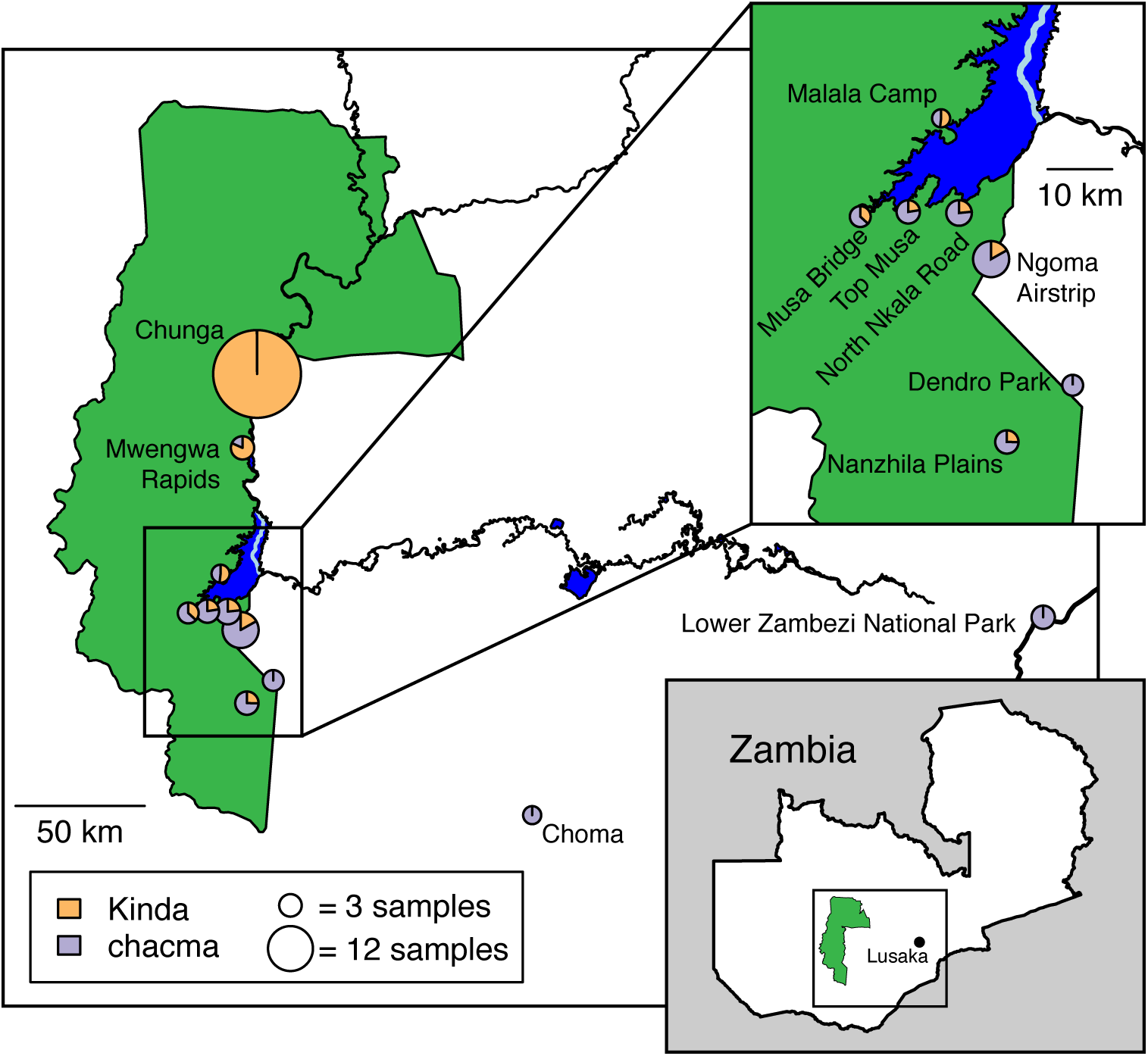
Average ADMIXTURE ancestry results (hybrid index) by locality. Circle areas are proportional to the number of individuals. The former course of the Kafue River prior to construction of the Itezhi Tezhi Dam between 1972 and 1976 is shown in light blue.

The midpoint of the cline, or the point where genetic ancestry from Kinda and chacma baboons is roughly equivalent, corresponded most closely to Malala Camp, an unmanned scout camp located near the confluence of the Chibila River and Lake Itezhi Tezhi (Figure 2). Baboons at this location are phenotypically intermediate in size and pelage, but resemble more closely Kinda baboons in overall appearance (Figure S8). South of Lake Itezhi Tezhi, groups from four localities in a narrow west-east band were on average mostly chacma, but contained a range of individuals from those with pure or mostly pure chacma baboon ancestry to those with more evenly mixed ancestry. Further south, at the Dendro Park private reserve, individuals were exclusively pure chacma baboons. Less than 15 km southwest in the Nanzhila Plains, while a small majority of individuals were pure chacma baboons, the remainder were roughly evenly mixed. See Figure S9 for hybrid indices by locality and Figure 2 for a map of localities.

We compared ancestry results from our autosomal marker set to ancestry results from mitochondrial and Y-chromosome DNA using nonparametric Wilcoxon signed rank tests (Table 2). We found that our ancestry results from autosomal markers were not significantly different from corresponding results from mitochondrial DNA. Ancestry results from Y-chromosome markers, however, were significantly different from ancestry results from both autosomal and mitochondrial DNA.

**Table 2:** Comparison of ancestry results from autosomal, mitochondrial, and Y-chromosome marker sets using nonparametric Wilcoxon signed rank tests. Results are shown for both the real sampling locales with data from at least one marker set and the expanded set of locales that included an additional 40 sampled from the geographic background.

| | Sampling locales | $W$ | $p$-value |
| --- | --- | --- | --- |
| autosomes vs. mitochondrial DNA | 40 real + 40 simulated | 1088 | 0.0725 |
|  | 40 real | 311 | 0.5919 |
| autosomes vs. Y chromosome | 40 real + 40 simulated | 1882 | < 0.0001 |
|  | 40 real | 630 | < 0.0001 |
| mitochondrial DNA vs. Y chromosome | 40 real + 40 simulated | 1350 | < 0.0001 |
|  | 40 real | 495 | < 0.0001 |

### Signatures of extreme introgression

In order to identify genes with extreme patterns of introgression, we used the Bayesian genomic cline model of Gompert and Buerkle (2011) implemented in bgc (Gompert & Buerkle 2012). We defined genes as having excess patterns of introgression if their posterior 95% credible intervals for *α*_*i*_ or *β*_*i*_ were entirely greater than or less than 0. From a total of 2,249 genes in our dataset, we identified 3 genes with extremely negative *α*_*i*_, 52 genes with extremely positive *α*_*i*_, 1 gene with extremely negative *β*_*i*_, and 5 genes with extremely positive *β*_*i*_ (Table S2, Table S3, and Figure S12). Positive *α*_*i*_ indicates an excess probability of chacma baboon ancestry while negative *α*_*i*_ indicates an excess probability of Kinda baboon ancestry relative to the genome-wide hybrid index. The high discrepancy between the number of genes with negative and positive *α*_*i*_ possibly indicates a higher rate of introgression of chacma baboon gene variants, but may also be a methodological artifact. Comparison between hybrid indices inferred by ADMIXTURE and bgc revealed a strong correlation, but the slope indicated that hybrid indices estimated by bgc were on average lower than corresponding estimates by ADMIXTURE (Figure S13). In contrast to the unsupervised approach in ADMIXTURE, bgc requires prior specification of parental populations. Because the *α*_*i*_ parameter is estimated based on a comparison between the per-locus probability of ancestry and the genome-wide hybrid index estimated by bgc, an inaccurately low hybrid index may artificially inflate the number of loci with high per-locus probabilities of chacma baboon ancestry. The lower sample size (*n* = 20) combined with the possibility of low levels of admixture in the chacma baboon parental sample may explain the systematic underestimation of chacma baboon ancestry by bgc.

While the above caveat applies to genes with positive *α*_*i*_, it conversely gives greater credence to genes identified as having extremely negative *α*, indicating excess Kinda baboon ancestry. These genes were *AMPH, KMT2E*, and *LY96*. It also does not affect estimates of *β*_*i*_, which is not influenced by the direction of introgression. Positive values of *β*_*i*_ indicate narrow clines and suggest potential roles in barriers to reproduction (Gompert & Buerkle 2011). The genes identified were *AACS, LIMK1, LY96, TMEFF2*, and *TMEM178*. Negative values of *β*_*i*_ indicate wide clines and suggest potential roles in adaptive introgression (Gompert & Buerkle 2011). *ODF2* was the only gene identified in this category.

We assessed the concordance of the bgc model with biological expectations by comparing cline parameter values of genic and nongenic SNPs. We predicted that genic SNPs, which are more likely to be functional, would be associated with stronger barriers to reproduction in the hybrid zone (Hvala *et al.* 2018) and would thereby have higher *β*_*i*_ values. We found statistically significant support for this prediction (Wilcoxon rank sum test, *p* = 0.0443).

### Gene function and biological pathway analysis

We conducted enrichment analyses in order to identify Gene Ontology (GO) terms (Gene Ontology Consortium 2000, 2015) and PANTHER pathways (Mi *et al.* 2013b; a) associated with extreme overall shifts in the genomic cline parameters *α*_*i*_ and *β*_*i*_. We identified 12 GO terms enriched for positive *α*_*i*_, 16 GO terms enriched for negative *α*_*i*_, 12 GO terms enriched for positive *β*_*i*_, and 14 GO terms enriched for negative *β*_*i*_ (Table S4). We also identified 4 PANTHER pathways enriched for positive *α*_*i*_, 1 PANTHER pathway enriched for positive *β*_*i*_, and 1 PANTHER pathway enriched for negative *β*_*i*_ (Table 3). We did not identify any pathways enriched for negative *α*_*i*_. Corrections for multiple comparisons were not applied for these exploratory analyses.

**Table 3:**
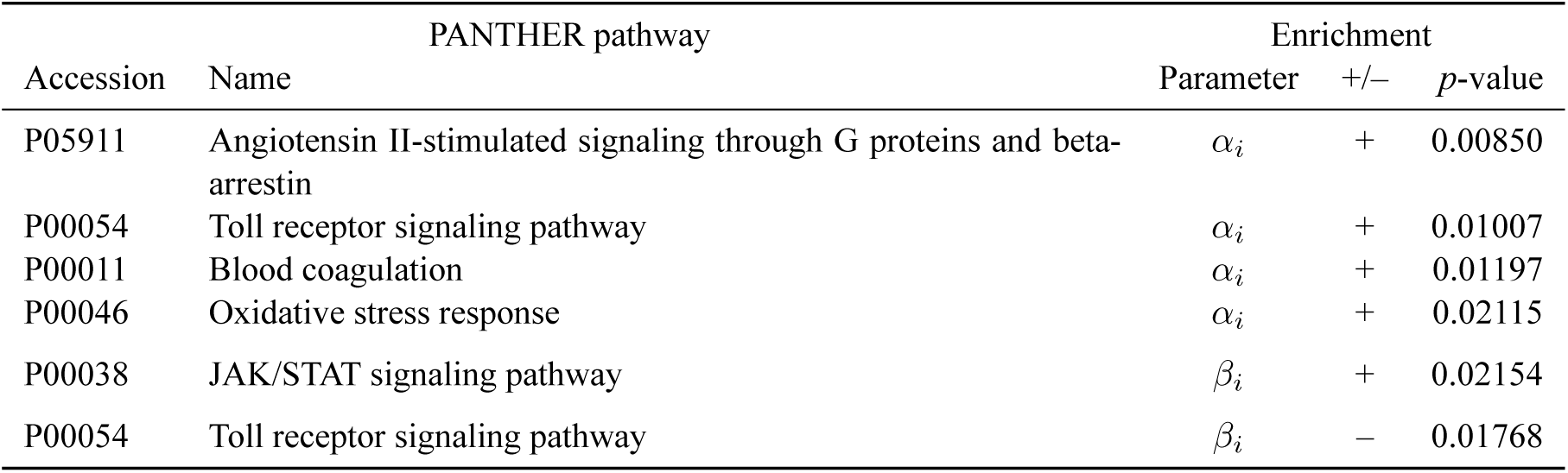
PANTHER pathways with significantly enriched *α*_*i*_ or *β*_*i*_ cline parameters. Only pathways with *p* < 0.025 for any of four one-tailed enrichment tests (positive *α*_*i*_, negative *α*_*i*_, positive *β*_*i*_, negative *β*_*i*_) are shown here. Key: +, test for positive parameter value; –, test for negative parameter value.

While differences between hybrid indices estimated by ADMIXTURE and bgc (Figure S13) affect the sign of *α*_*i*_ estimates and consequently the identification of candidate genes with extreme *α*_*i*_ as previously discussed, they do not affect the enrichment of *α*_*i*_ because enrichment analyses test for extreme parameter estimates relative to the full distribution of parameter values rather than 0.

## Discussion

### Geographic structure and extent of the hybrid zone

We found that Kinda and gray-footed chacma baboons hybridize over a wide area spanning at least 100 km in the Kafue River Valley in Zambia. These results corroborate phenotypic observations as well as previously reported mitochondrial-DNA and Y-chromosome results (Jolly *et al.* 2011).

Ansell (1978), in the last published survey of Kafue National Park baboons prior to our work (Jolly *et al.* 2011), remarked that specimens of phenotypically chacma baboons were collected as far north as the vicinity of Lubalunsuki Hill and Itumbi (approximately 15.50° S., 25.97° E.). Phenotypically Kinda baboons were observed in the area as well, but no indications of interbreeding were known. It now seems possible that hybridization was already occurring given the geographic proximity of the species and the extent of the hybrid zone today, but if these descriptions are otherwise accurate, then the hybrid zone has shifted considerably south in the past half-century. Baboons near Lubalunsuki Hill today resemble phenotypically Kinda baboons (Figure S14) and the center of the hybrid zone is 30 – 40 km south of Lubalunsuki. Furthermore, Kinda baboon mitochondrial haplotypes are found as far south as the Ngoma Airstrip, 55 km south of Lubalunsuki, while Y-chromosome haplotypes, as well as intermediate phenotypes, are found as far south as the Nanzhila Plains, 80 km south of Lubalunsuki. These findings suggest either that Ansell’s (1978) descriptions were inaccurate or that there has been strong directional movement of Kinda baboons (roughly 1-2 km per year), including females, into the chacma baboon distribution. This movement may have come from the north, but may also have come from the west, where there is an expanse of miombo forest corresponding to typical Kinda baboon habitat from which little information is available.

Between 1972 and 1976, construction of a new hydropower dam at Itezhi Tezhi flooded a large section of the Kafue River, as well as its upstream tributaries on the western bank, forming the artificial Lake Itezhi Tezhi (see Figure 2). The extent to which this anthropogenic event impacted baboon hybridization is unknown, but raises intriguing questions. The impacts of flooding on the local ecology might have created conditions more or less favorable to either species. The flooding would have also displaced existing groups of baboons, potentially precipitating contact and subsequent hybridization between species. Based on Ansell’s (1978) descriptions, it seems likely the lake flooded primarily chacma baboon territory and the cline of the hybrid zone, which today begins northwest of Lake Itezhi Tezhi and extends around and to the area south of the lake, formed in the time since. If a hybrid zone already existed, however, roughly coinciding with the present cline, then the formation of Lake Itezhi Tezhi bisected a portion of the hybrid zone, creating a barrier between the area west of Lake Itezhi Tezhi and the area immediately south of the Itezhi Tezhi Dam. A landscape genetic approach comparing baboons on either side of this barrier could help resolve this issue.

### Y-chromosome discordance

Our results indicate that there is a sizeable average excess of Kinda baboon Y-chromosome ancestry relative to not only mitochondrial DNA, as previously reported by Jolly *et al.* (2011), but the remainder of the nuclear genome as well (Figure 3). Interestingly, the unidirectional Y introgression relative to both mitochondrial DNA and the autosomes parallels findings in another papionin group, macaques (genus *Macaca*), in which the *M. mulatta* Y chromosome has apparently introgressed into *M. fascicularis* populations, reaching as far south as the Isthmus of Kra in mainland southeast Asia and resulting in discordance with both the *M. fascicularis* mitochondrial genome and phenotype (Tosi *et al.* 2002).

**Fig 3:**
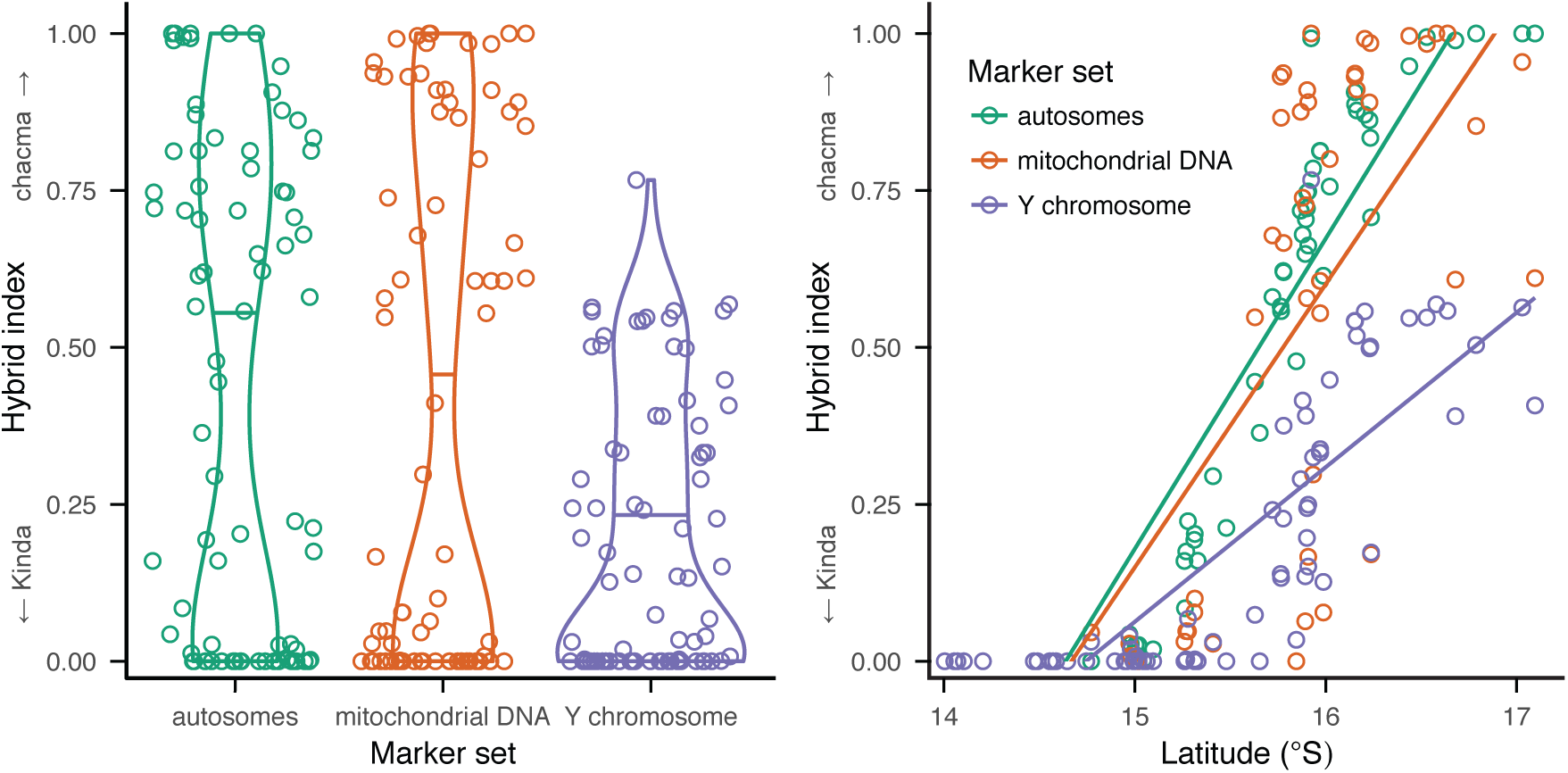
A comparison of ancestry results between autosomal, mitochondrial, and Y-chromosome marker sets. Ancestry estimates from each marker set were averaged for each site and interpolated over the study area to facilitate comparison among all sample points (Figure S10 and Figure S11). Violin plots in the left pane show the distribution of hybrid indices estimated from the three marker sets, with mean estimates indicated by horizontal lines. The scatter plot in the right pane shows hybrid indices plotted against latitude. Mitochondrial and Y-chromosome data are from Jolly et al. (2011).

As previously discussed by Jolly *et al.* (2011), mitochondrial/Y-chromosome discordance in Kafue National Park baboon hybrids suggests strong unidirectional Kinda baboon Y-chromosome introgression. In the absence of autosomal information, this is consistent with the common case in species in which males are the dispersing sex (Zinner *et al.* 2011) and indicates that the hybrid zone could be driven primarily by immigration of male Kinda baboons. A probable eventual outcome of this scenario is “nuclear swamping”, by which successive backcrossing between hybrids and the influx of foreign males largely replaces the native population’s nuclear genome but may retain the native population’s mitochondrial gene pool (Rohwer *et al.* 2001; Roca *et al.* 2004; Ting *et al.* 2008). Our results, however, reveal that the Kinda baboon Y chromosome has introgressed more extensively on average than autosomal DNA, indicating either that insufficient time has passed for nuclear swamping to occur or that migration of male Kinda baboons is not the principal factor driving the hybrid zone and other processes may be at work.

Jolly *et al.* (2011) proposed several hypotheses that explain strong unidirectional male Kinda baboon introgression without requiring unidirectional migration. One hypothesis is that reproductive pairings between male chacma baboons and female Kinda baboons are disadvantaged because of reproductive incompatibilities, perhaps associated with obstetric challenges that result from the unusually extreme size difference between potential mates (or more precisely, between mothers and their hybrid fetuses). Another hypothesis is that the diminutive and juvenescent appearance of adult Kinda baboons leads to impaired recognition of potential female mates and/or male competitors by male chacma baboons. The latter effect may be compounded by the unusual grooming behavior in Kinda baboons by which males actively initiate and maintain grooming associations with females even when females are not in estrus (Weyher *et al.* 2014). Both of these hypotheses are consistent with our results, as they suggest that while male chacma baboons are disadvantaged in the hybrid zone, subsequent backcrossing between male hybrid and female chacma baboons may not be similarly inhibited. Such backcrossing would counteract replacement of the chacma baboon nuclear genome while explaining the extreme introgression of the Kinda baboon Y chromosome.

A neutral model, however, based on sex differences in reproductive skew and the lower effective population size of the Y chromosome could also explain the unidirectional introgression of the Y chromosome. The uniparental inheritance of the Y chromosome, as well as the mitochondrial genome, results in reduced effective population sizes of both markers equaling 1/4 the effective population size of the autosomes assuming even sex ratios and the absence of sex-biased processes (Charlesworth 2009). If male baboons have higher variance in reproductive success than females, however, the variance effective population size of the patrilineally inherited Y chromosome would be even lower than that of the matrilineally inherited mitochondrial genome. Under these conditions, genetic drift could produce large changes in Y-chromosome frequencies relatively quickly. Given that primate males of many taxa including gray-footed chacma baboons (Bulger 1993) tend to have higher variance in reproductive success relative to females (Kutsukake & Nunn 2006), the neutral model could plausibly explain the unidirectional introgression of the Y chromosome.

### Non-neutral introgression

We used Bayesian genomic clines to identify putative loci experiencing enhanced, restricted, or directional introgression across the hybrid zone. These analyses identified several genes, biological processes, and biological pathways with signals of non-neutral introgression. Importantly, this analysis does not represent a comprehensive scan of the genome. Our use of double-digest RADseq, while effective for optimizing our read coverage and cost of sequencing (Peterson *et al.* 2012), also limited the fraction of the genome available for this analysis. Given that our average distance between loci likely exceeds the extent of linkage disequilibrium (Lowry *et al.* 2017), we expect to have missed a large fraction of genes, which also decreases the statistic power of subsequent enrichment analyses. Nevertheless, we were able to identify several intriguing patterns, described in subsequent sections.

### JAK/STAT signaling pathway

The Janus kinase (JAK) and signal transducer and activation of transcription (STAT) signaling pathway was the only biological pathway in our analysis enriched for positive *β*_*i*_ (Table 3), indicating an overall restricted pattern of introgression among components of this pathway. The JAK/STAT pathway functions as a signaling mechanism for a variety of systems and underlies diverse responses such as mammary gland development, hematopoiesis, and immune cell development (Watson & Burdon 1996; Ward *et al.* 2000; Ghoreschi *et al.* 2009). Intriguingly, it also underlies pathways related to organism growth, for example by serving as the signaling mechanism for the growth hormone (GH) and insulin-like growth factor 1 (IGF1) axis that plays an important role in organism growth through chondrocyte proliferation and differentiation (Nilsson *et al.* 1994; Vottero *et al.* 2013). JAK/STAT signaling has also been shown to play a role in leptin-induced chondrocyte differentiation, which is mediated through STAT3 (Ben-Eliezer *et al.* 2007).

Notably, the JAK/STAT signaling pathway has been found to be associated with high *F*_*ST*_ between pure Kinda and chacma baboons, suggesting directional selection in one or both species (Chiou *et al.* 2019). The enrichment of JAK/STAT signaling for positive *β*_*i*_ in the present study suggests that it may also underlie isolating mechanisms in these species. The relationship of JAK/STAT components to body size differences in baboons has not yet been confirmed or characterized. If changes in the JAK/STAT pathway are indeed related to the extreme differences in body size between Kinda and chacma baboons, our results suggest that these differences may be responsible for decreased fitness of hybrid offspring and that the specific genes under selection include components of the JAK/STAT signaling pathway. As suggested by Jolly *et al.* (2011), the selection may be strongest against reproduction between male chacma-like baboons and female Kinda-like baboons, possibly due to obstetric limitations differentially impacting hybrid offspring of small Kinda-like mothers or issues with mate or competitor recognition due to the juvenescent appearance of both female and male Kinda baboons. It may also, however, affect reproduction between male Kinda-like baboons and female chacma-like baboons, whose body size dimorphism would be on the low extreme of that found between mating partners in extant papionin monkeys (Delson *et al.* 2000). Barriers to reproduction are possible in both reciprocal cases but our findings at present are unable to distinguish between the two.

Alternatively, restricted introgression within this pathway may reflect epistatic selection acting to maintain concerted functions by conserving interactions among multiple components of this pathway in the presence of introgressive gene flow (Gavrilets 1997). Given its diverse functions and its involvement as a signaling pathway for a variety of cytokines and growth factors (Harrison 2012), this hypothesis at present is more parsimonious.

Within the JAK/STAT signaling pathway, the protein inhibitor of activated STAT (PIAS) genes *PIAS1* and *PIAS4*, as well as *STAT2*, were associated with the highest *β*_*i*_ values, although none was identified individually as a candidate based on posterior 95% credible intervals (Table S5). PIAS1 has been shown to inhibit STAT1 (Liu *et al.* 1998), which has anti-proliferative effects in cells including chondrocytes (Sahni *et al.* 1999). PIAS1 has also been shown to be a regulator of SOX9, a transcription factor that plays critical roles in developmental processes including chondrogenesis and testis determination (Oh *et al.* 2007).

### Immune adaptation in the hybrid zone

The toll-like receptor (TLR) signaling pathway was the only biological pathway in our analysis that was enriched for negative *β*_*i*_ (Table 3), indicating that its cline was wider than average. TLR signaling is involved in the recognition of pathogenic microbes and plays a role in the activation of both innate immunity and antigen-specific acquired immunity (Akira & Takeda 2004).

Enrichment for negative *β*_*i*_ in this pathway may reflect adaptive introgression related to the introduction of novel pathogenic defenses. Notably, similar adaptive introgression of immune system components from archaic humans has significantly shaped the immune systems of modern human populations (Abi-Rached *et al.* 2011; Dannemann *et al.* 2016; Enard & Petrov 2018). For instance, haplotypes in modern humans obtained through archaic human introgression include three TLRs (*TLR6*-*TLR1*-*TLR10*) demonstrably associated with microbial resistance (Dannemann *et al.* 2016). An analysis of longer, and likely adaptive, segments of Neanderthal ancestry in European genomes has also found an enrichment of proteins that interact with viruses, particularly RNA viruses (Enard & Petrov 2018). Also in Europeans, admixture with Neanderthals has introduced functional variants that differentially regulate the response to TLR7/8 stimulation at at least one locus (Quach *et al.* 2016). In our analysis, the TLR signaling pathway was also enriched for positive *α*_*i*_ values, indicating that introgression of alleles within this pathway primarily took place from chacma into Kinda baboon populations.

Two genes in the TLR signaling pathway were included in our analysis: *MAP2K2* and *NFKBIE* (Table S6), both of which encode kinases that play integral roles in the downstream signaling cascade (Cohen 2014).

Interestingly, the blood coagulation pathway was also enriched with positive *α*_*i*_ values, suggesting excess chacma baboon ancestry. The coagulation pathway overlaps with the innate immune response and has been a common target of positive selection throughout vertebrate evolution (Rallapalli *et al.* 2014). Neanderthal alleles in modern Europeans have also been associated with phenotypes including coagulation, suggesting that they may have undergone adaptive introgression (Simonti *et al.* 2016).

At the individual gene level, we also identified five candidate genes including *LY96* that had a wide cline (negative *β*_*i*_) relative to the genome-wide average. LY96, also known as MD2, associates with TLR4 and confers responsiveness to lipopolysaccharide (LPS) (Shimazu *et al.* 1999), thus providing another link to TLR signaling (LY96 is not a component of TLR signaling in PANTHER). *LY96* was also identified as having a negative *α*_*i*_, indicating excess Kinda baboon ancestry. Notably *LY96* exhibits the opposite directional pattern from the PANTHER TLR pathway, indicating that introgression of different immune components may proceed in both directions.

Our Gene Ontology results (Table S4) were consistent with the view that immune-related variants exhibit enhanced introgression. Wide clines (negative *β*_*i*_) were enriched with terms including “negative regulation of viral process” and “immune system development”. Excess chacma ancestry (positive *β*_*i*_) was enriched with terms such as “response to antibiotic” while excess Kinda ancestry (negative *β*_*i*_) was enriched with terms such as “regulation of T cell differentiation”, “T cell differentiation”, and “regulation of lymphocyte differentiation”. Narrow clines (positive *β*_*i*_) were not enriched with any terms having an obvious relation to immune function.

### Outer dense fiber protein 2

*ODF2* (outer dense fiber protein 2) in our study was the only gene identified as having extremely negative *β*_*i*_, indicating a wide cline, as well as as extremely positive *α*_*i*_, indicating excess chacma baboon ancestry. These results suggest that *ODF2* is a candidate gene for adaptive introgression, with the direction of introgression primarily taking place from chacma baboon populations into Kinda baboon populations. This result was driven by a single SNP (chr15:9561907) located in an intronic region of *ODF2*.

Outer dense fibers are a class of cytoskeletal structures that are specific to sperm tails (Brohmann *et al.* 1997; Hoyer-Fender *et al.* 1998; Soung *et al.* 2006). ODF2 has been shown to be critical for the structural integrity of the sperm flagellum. Inhibition of tyrosine phosphorylation of ODF2 adversely affects sperm motility during capacitation in hamsters (Mariappa *et al.* 2010). In ODF2-knockout mouse lines, males with high percentages of chimerism exhibit altered sperm tail structure and function, with some sperm exhibiting additional defects including bent tails displaying abnormal motility (Tarnasky *et al.* 2010).

*ODF2* has been subject to recent positive selection in European human populations (Voight *et al.* 2006). Intriguingly, it has also been associated with substantial structural differences in chimpanzee DNA as part of a suite of genes showing markedly chimpanzee-specific changes relative to humans and orangutans (Kim *et al.* 2011). Chimpanzee females exhibit polyandrous mating and, as would be predicted, chimpanzee males exhibit an extremely large relative testis size compared to humans, gorillas, and orangutans (Harcourt *et al.* 1981). Female polyandry is predicted to promote not only the evolution of larger testes and thereby ejaculate quantities, but also higher-quality (e.g., longer tails and faster) sperm (Fitzpatrick *et al.* 2009; Schmera *et al.* 2016).

Adaptive introgression of *ODF2* is surprising at face value given that sperm traits are commonly subject to dysgenesis in the context of an alien genome (e.g., Dobzhansky 1934). In *Lepomis* sunfish for instance, hybrid sperm are fertile in the absence of competition but are outcompeted in the presence of sperm from either parental species (Immler *et al.* 2011). Conspecific sperm precedence is also observed in competition between bluegill and pumpkinseed sunfish sperm for pumpkinseed sunfish eggs, but not in the reciprocal case, potentially explaining unidirectional hybridization in this system (Immler *et al.* 2011). In the European house mouse hybrid zone, hybrid mice exhibit significantly reduced sperm count and sperm velocity, indicating widespread hybrid dysgenesis of sperm traits (Turner *et al.* 2011; Albrechtová *et al.* 2012).

Given the costs of sperm production (Wedell *et al.* 2002), selection on sperm quantity and quality is expected to be particularly important in taxa with higher rates of polyandrous mating (Ginsberg & Huck 1989). In baboons and other animals, the priority-of-access model (Altmann 1962) posits that male access to estrous females is determined according to rank. Throughout the estrous period, males engage in mate-guarding in order to monopolize mating access and ensure paternity. This system may be subverted, however, by mechanisms including male coalitionary behavior, sneak copulations, and female choice (Alberts *et al.* 2003). Gray-footed chacma baboons have previously been shown to conform well to the priority-of-access model and exhibit one of the strongest correlations between rank and mating among baboons (Bulger 1993). While information on Kinda baboons is scarce, emerging information suggests a different picture. Male Kinda baboons notably engage at high frequency in grooming associations with females in which they play an active role in initiating and maintaining the associations even when females are pregnant, lactating, or otherwise not in estrus (Weyher *et al.* 2014). Preliminary data from a Kinda baboon study population at Kasanka National Park show a relatively high frequency of matings not predicted by dominance hierarchies and demonstrate that polyandrous mating by females occurs at least some of the time (A. Weyher, pers. comm.).

These characteristics would seem to imply a heightened role of sperm competition in Kinda baboons relative to chacma baboons. This prediction is further supported by morphometric analyses demonstrating that male Kinda baboons have larger testes, scaled to body size, relative to all other evaluated congeneric species except anubis baboons (Jolly et al., in review), corroborating parallel investigations of relative testis sizes in hamadryas baboons, which are monandrous, and anubis baboons, which are both polygynous and polyandrous (Jolly & Phillips-Conroy 2003, see also 2006).

Our results, however, suggest that the chacma baboon *ODF2* variant traverses the species boundary to a greater extent than the Kinda baboon variant, contrary to our expectations. Comparison to the European house mouse hybrid zone (Albrechtová *et al.* 2012) offers an intriguing explanation. Like the present baboon hybrid zone (Jolly *et al.* 2011), the *Mus musculus musculus* Y chromosome has introgressed into *Mus musculus domesticus* populations in apparent disregard of Haldane’s rule. While Albrechtová *et al.* (2012) found that hybrids overall exhibited decreased sperm counts, this effect was more than rescued in apparently *domesticus* males by the presence of the invading *musculus* Y chromosome. This finding is surprising given that respective *domesticus* and *musculus* sperm traits and Y chromosomes evolved in concert subject to natural selection in the parental mouse populations, and would likely encounter Dobzhansky-Muller disadvantages when placed in a novel genetic background. The effect of the *domesticus* Y chromosome on sperm traits instead implies that there is a sperm-related advantage in the presence of the invading Y chromosome that sufficiently balances and even overcomes this effect.

The present hybrid zone, where the Kinda baboon Y chromosome experiences unidirectional introgression but the chacma baboon sperm-related allele experiences disproportionate success, presents a compelling parallel to the house mouse system (Albrechtová *et al.* 2012). The house mouse analogy suggests that hybrid dysgenesis of sperm-related traits may decrease reproductive fitness in the Kinda and chacma baboon species boundary in the Kafue River valley due to Dobzhansky-Muller incompatibilities. This effect, however, may be mitigated or even overcome by beneficial interactions with the introgressed Y chromosome of one parental species, in this case the Y chromosome of the Kinda baboon. This hypothesis could not be tested in the present study due to the challenges of genotyping Y-chromosome markers from our dataset and the lack of phenotypic data on sperm characteristics, but offers an intriguing alternative explanation for the surprising unidirectional introgression of the Kinda baboon Y chromosome.

## Acknowledgments

We thank the Zambia Wildlife Authority (now the Department of National Parks & Wildlife) and the University of Zambia for granting permission and providing support for fieldwork. We also thank Anna Weyher for sharing unpublished information about Kinda baboon behavior. This study was funded by the National Science Foundation (BCS 1341018, SMA 1338524, BCS 1029302, BCS 1029323, BCS 1029451), the Leakey Foundation, and the National Geographic Society. The Genome Technology Center at NYU is supported by NIH/NCATS UL1 TR00038 and NIH/NCI P30 CA016087. K.L.C. is supported by NIH/NIA T32 AG000057.

## Data Accessibility

Sequence reads are deposited in the NCBI Sequence Read Archive (SRA) under BioProject PRJNA486659, with SRA accession numbers SRR7717274-SRR7717402. All code for this project is available on GitHub at https://github.com/kchiou/kafue-baboons-ddrad.

## Author Contributions

K.L.C., C.M.B., A.S.B., T.R.D., J.R., C.J.J., and J.P.C. designed the research and collected samples. K.L.C. performed the research. K.L.C. and C.M.B. analyzed the data. K.L.C. wrote the paper. All authors read, revised, and approved the manuscript.

