## Supplemental Figures and Tables for "Genome-wide ancestry and introgression in a Zambian baboon hybrid zone"

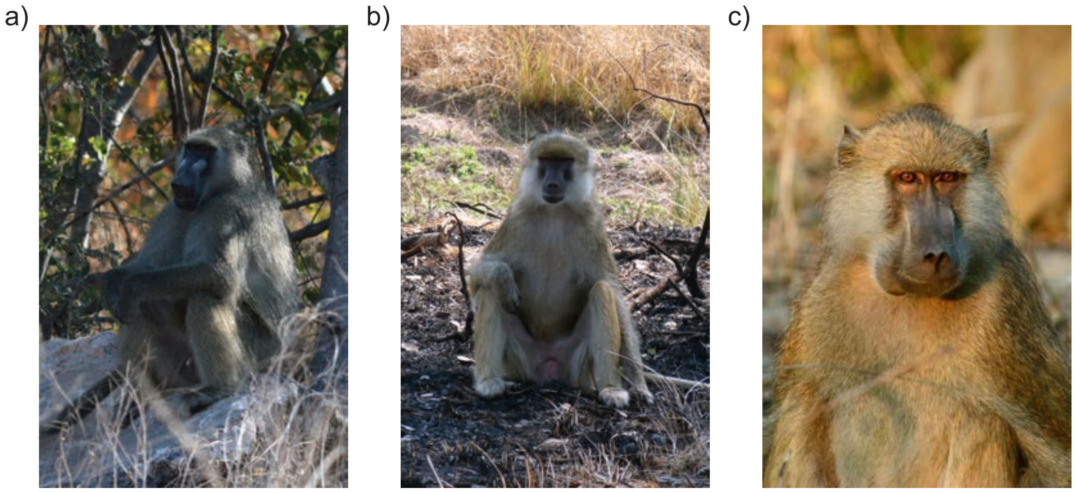


Figure S1: Photos of male a) gray-footed chacma, b) Kinda, and c) hybrid baboons. Photos are by Kenneth Chiou and licensed under Creative Commons (CC-BY-SA-4.0).

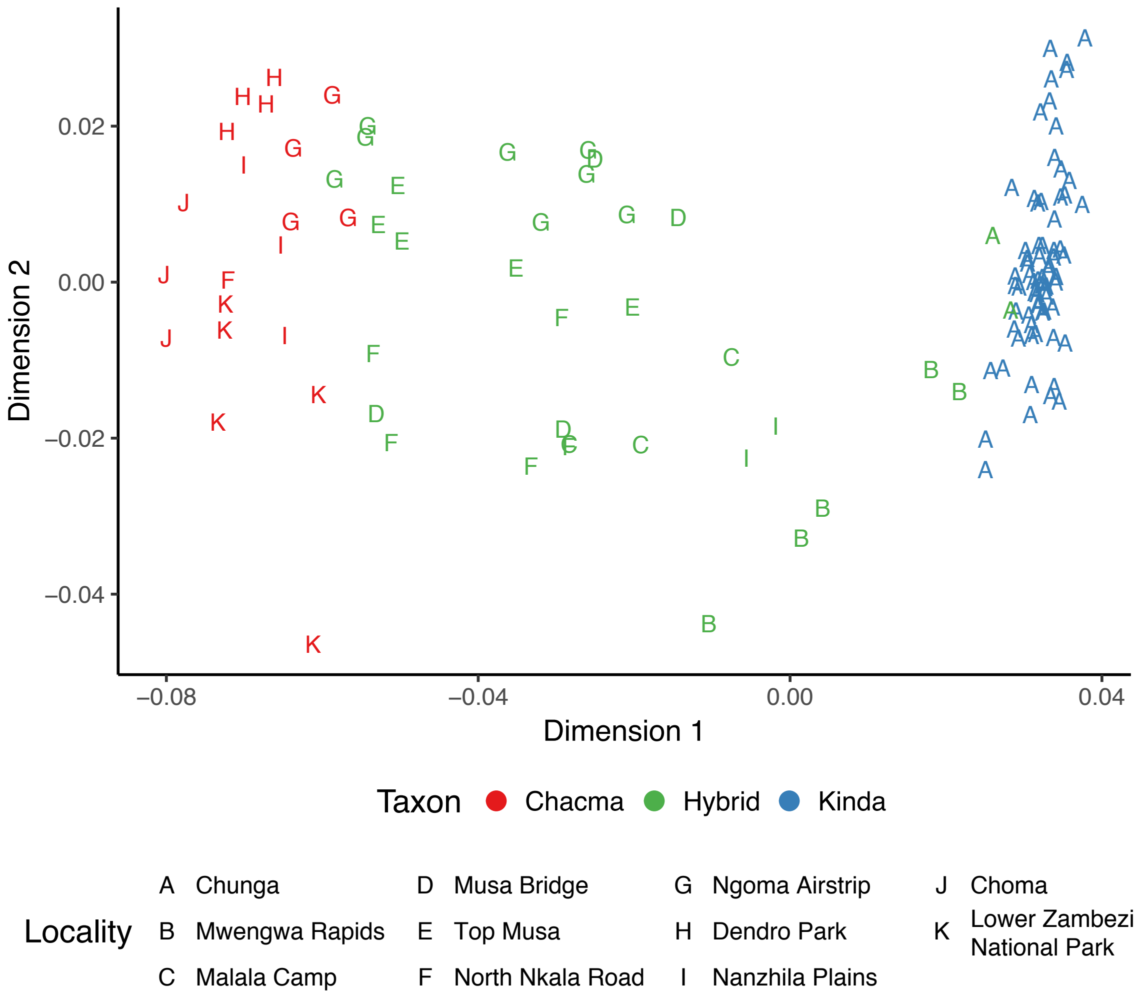


Figure S2: Multidimensional scaling of identity-by-state: first two dimensions. Points are colored based on their taxonomy inferred using ADMIXTURE (see Figure S5).

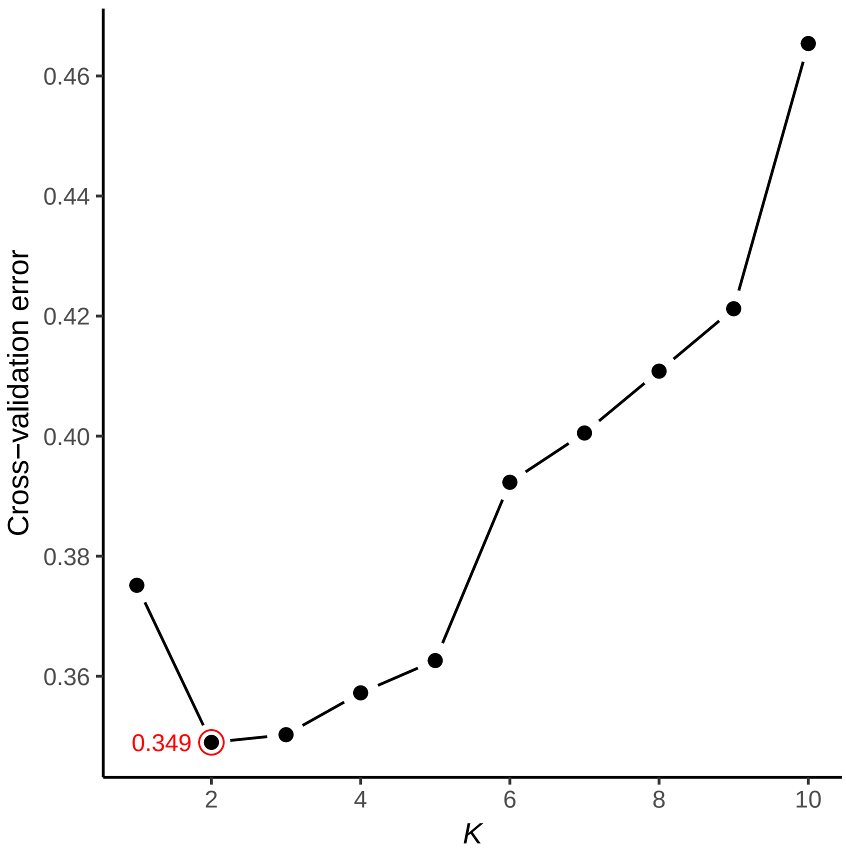


Figure S3: Cross-validated error results from ADMIXTURE runs in which $K$ varied from 1 to 10.

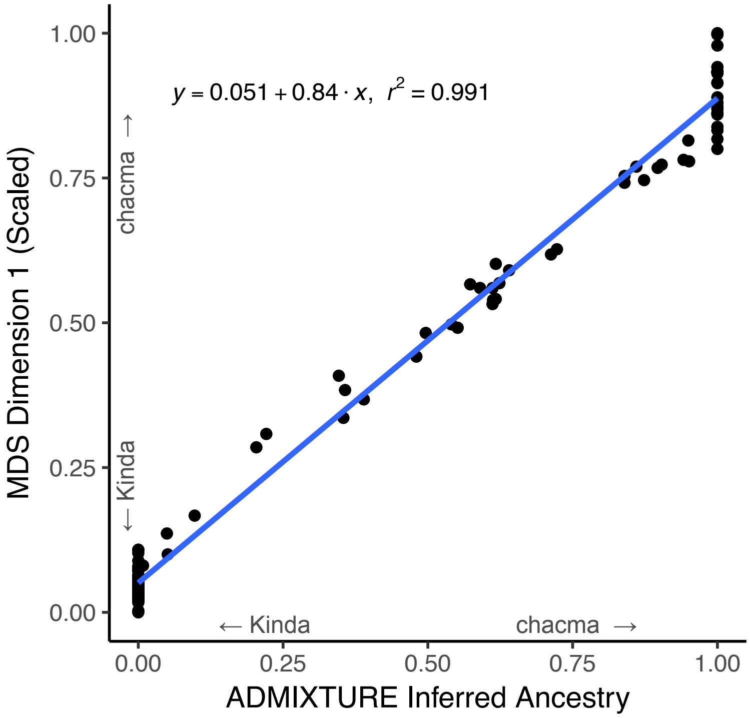


Figure S4: Concordance between multidimensional scaling and ADMIXTURE analysis. The first dimension of the MDS was first scaled from 0 to 1.

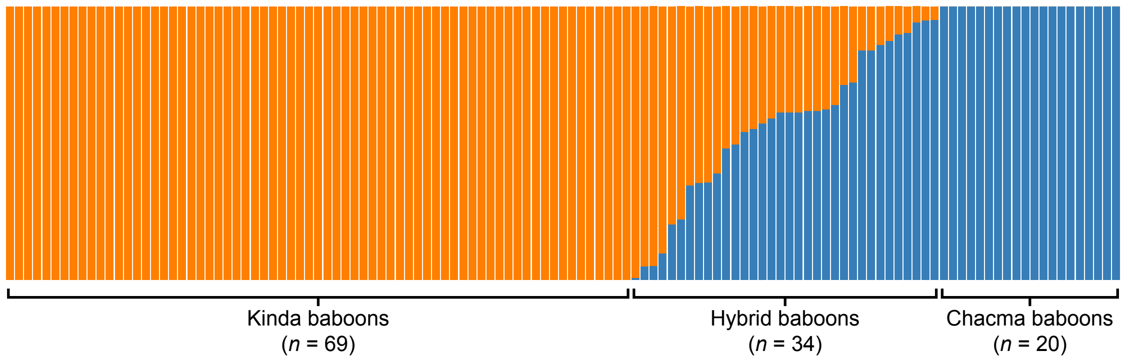


Figure S5: Taxonomic assignment based on ancestry estimates calculated using ADMIXTURE.

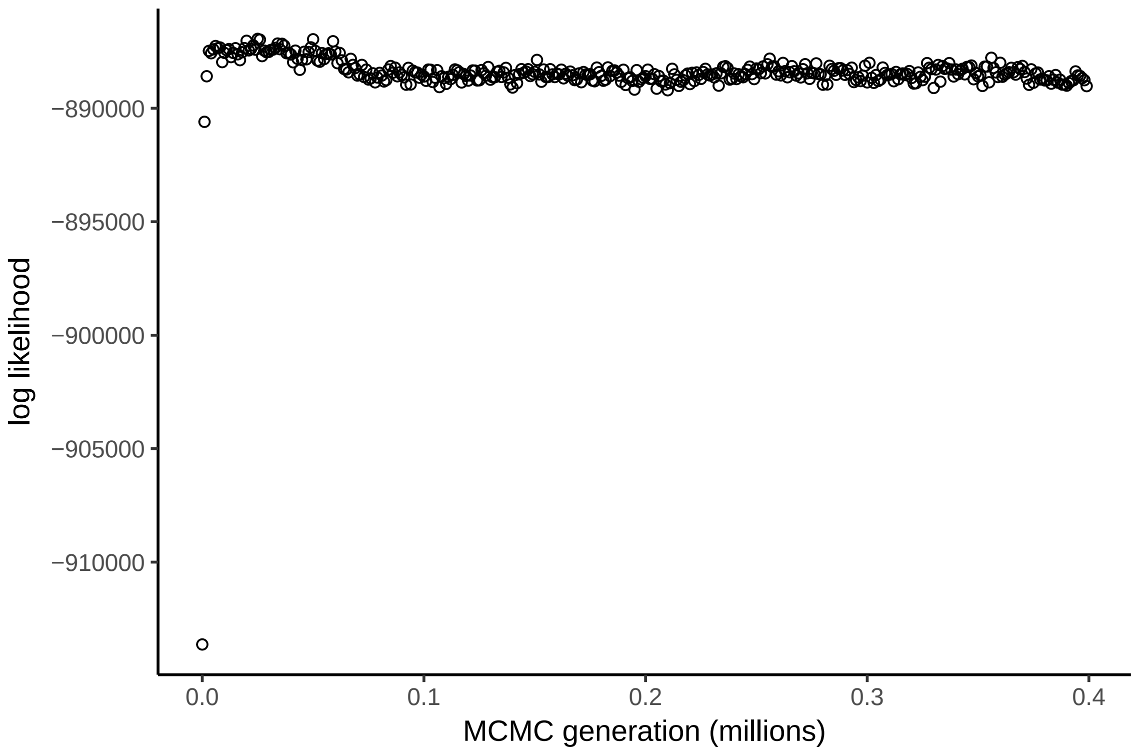


Figure S6: A representative example of a converged complete bgc chain of 400,000 generations. For this plot, chains are shown every 1,000 generations.

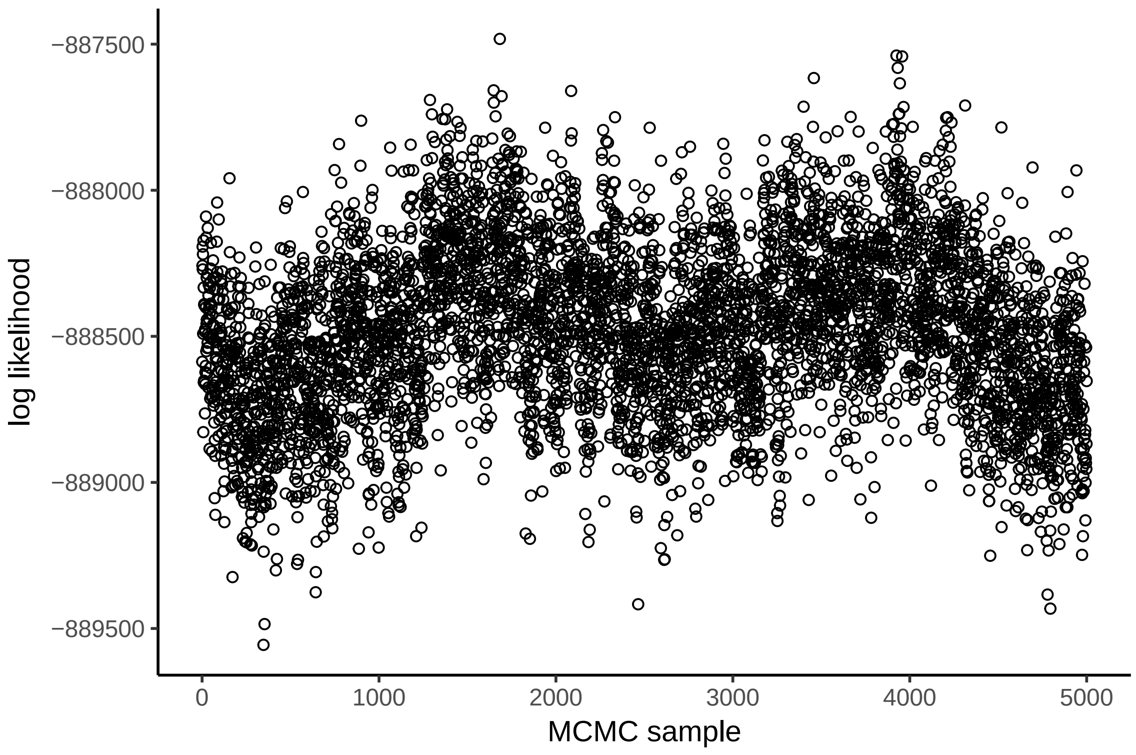


Figure S7: A representative example of a converged posterior BGC chain of 5,000 samples. This chain represents the last 200,000 generations of the chain shown in Figure S6, sampled every 40 generations.

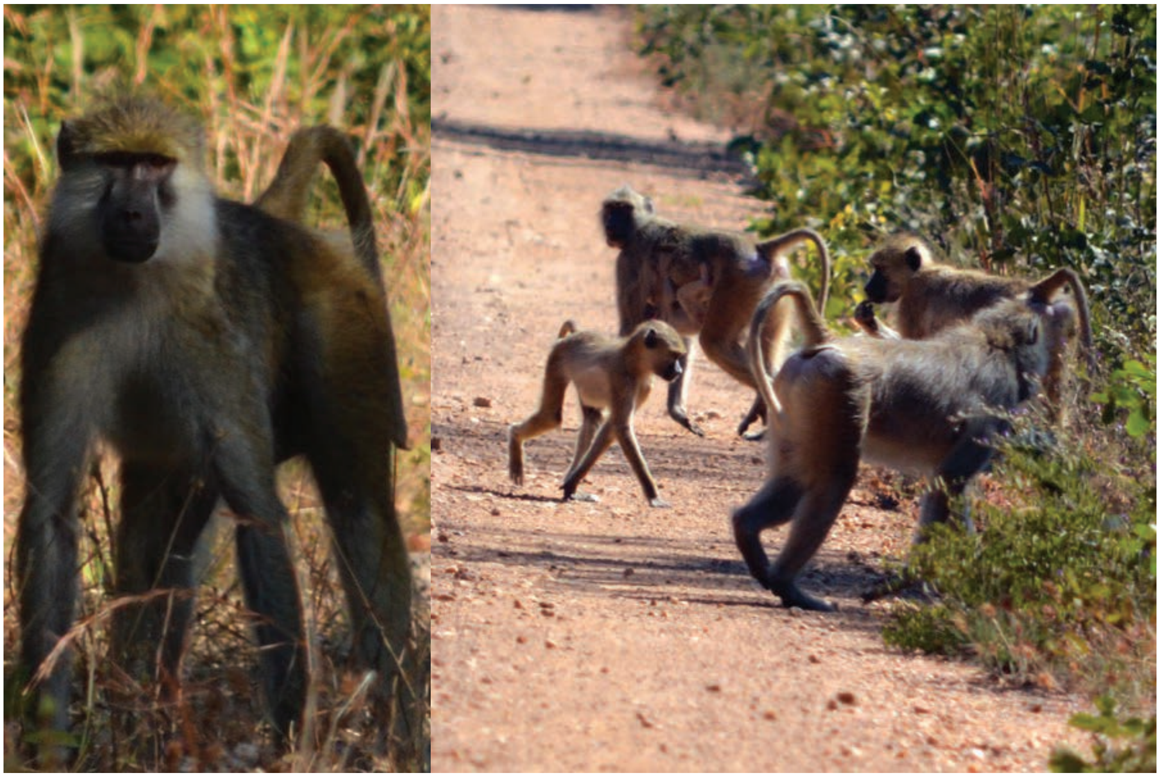

Figure S8: Baboons from Malala Camp resemble Kinda baboons superficially, with pink circumorbital skin, strongly contrasting cheek color, and a lanky appearance, but are intermediate in size and darker in coloration. In the right pane, the male in the foreground superficially resembles a gray-footed chacma baboon. Photos by Kenneth Chiou.

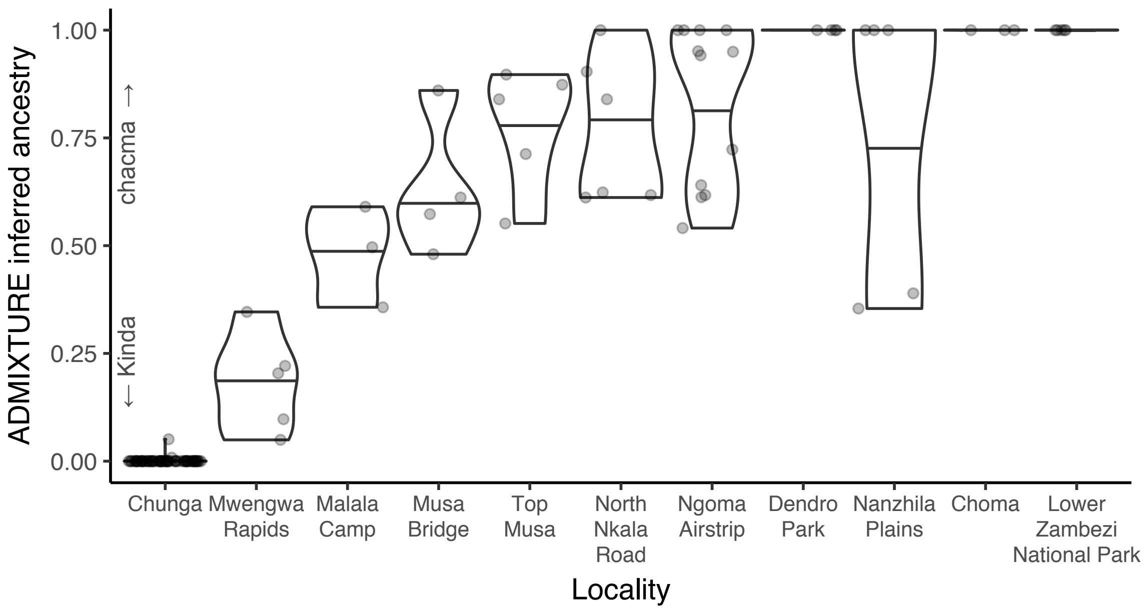


Figure S9: ADMIXTURE ancestry estimation results by locality.

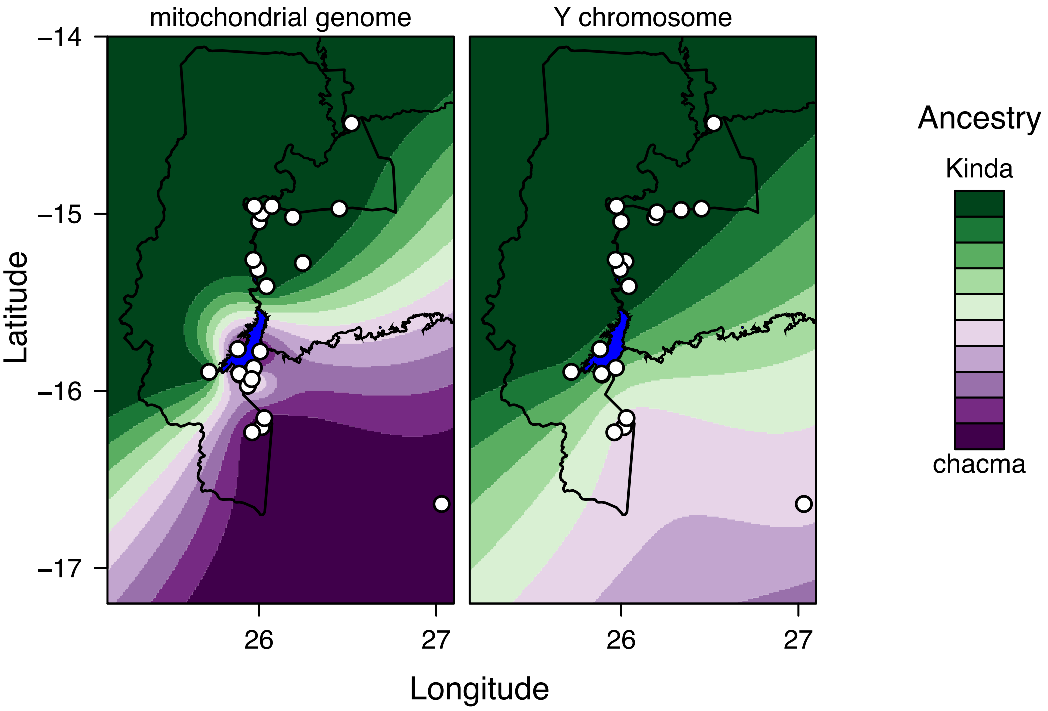


Figure S10: Geospatial interpolation of mean ancestry estimates for mitochondrial DNA and the Y chromosome. Mitochondrial and Y genotypes from Jolly et al. (2011) were averaged for each collection site and interpolated over the study area at 1 $km^2$ resolution using a Kriging surface model procedure.

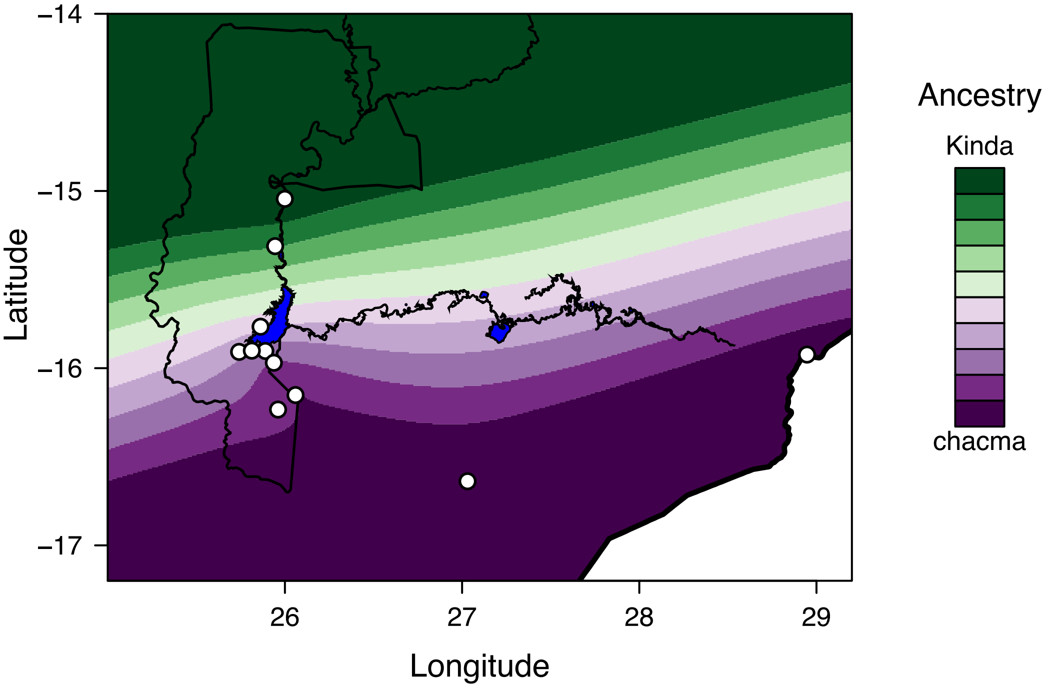


Figure S11: Geospatial interpolation of ancestry estimates for autosomal SNPs genotyped by ddRADseq. Ancestry estimates obtained from ADMIXTURE analysis (Figure 1) were averaged for each collection site and interpolated over the study area at 1 km$^2$ resolution using a Kriging surface model procedure. The geographic limit of this figure has been extended to the east in order to include the chacma baboon sites of Choma and Lower Zambezi National Park. As a consequence of this, ancestry estimates over much of the eastern half of this map are extrapolated and should be interpreted with caution, as they are almost certainly inaccurate.

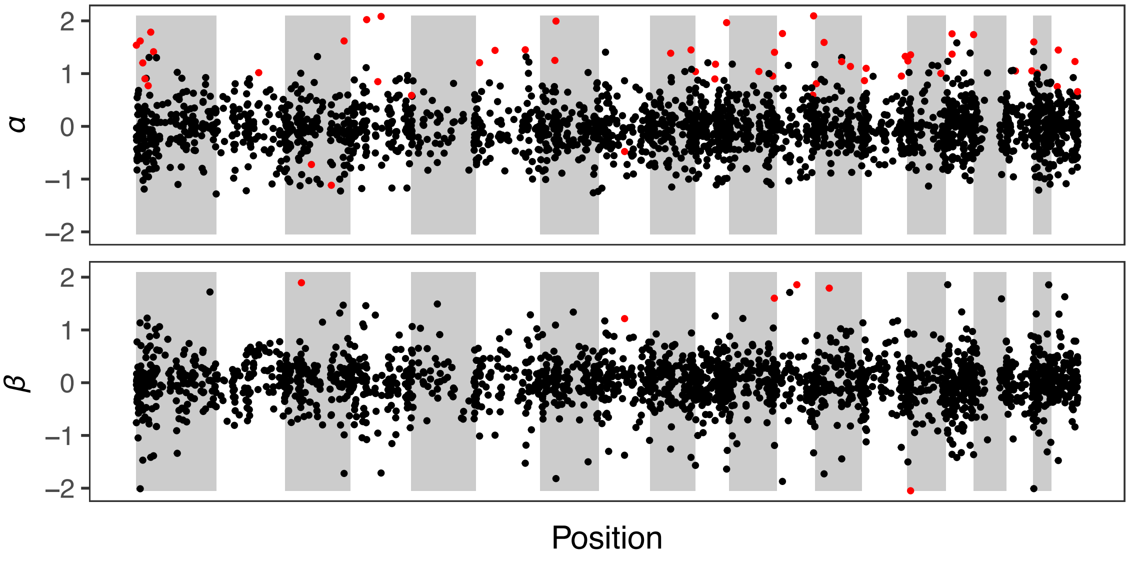


Figure S12: Distribution of $\alpha_i$ and $\beta_i$ estimates for protein-coding genes. Genes identified as extreme for a parameter by 95% Bayesian credible intervals are displayed in red. Background shading indicates the position of the autosomal chromosomes in order.

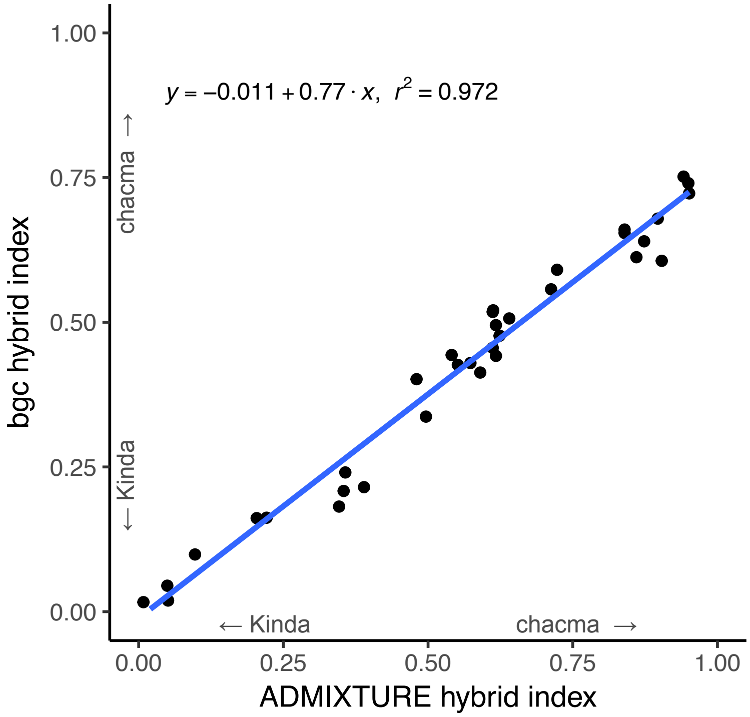


Figure S13: Concordance between hybrid indices estimated using ADMIXTURE and bgc.

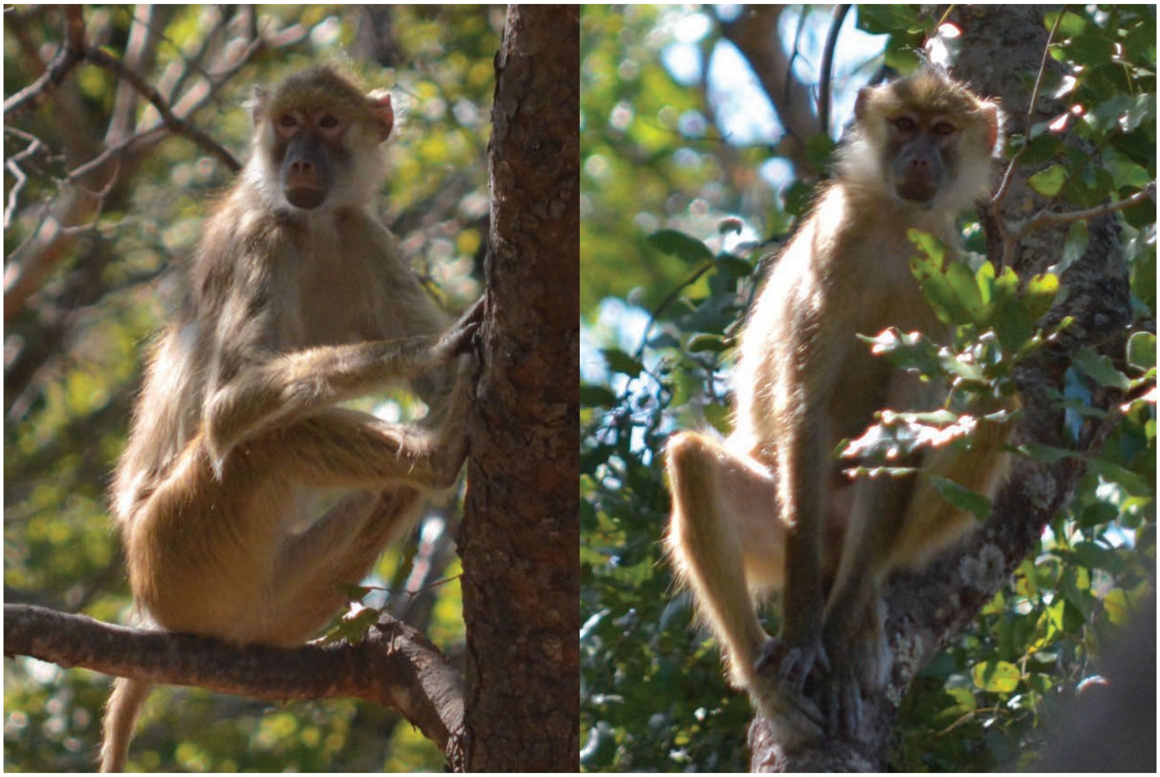

Figure S14: Baboons from Lubalunsuki Hill in the present day resemble Kinda baboons. Photos by Kenneth Chiou.

Table S1: Full list of samples included in this analysis. Samples that were sequenced but failed quality-control filters are not included in this table. Hybrid indices and ancestry determinations are based on ADMIXTURE analysis. Geographic coordinates for all localities are given in Table 1.

| Sample ID | Sample type | Locality | Hybrid index | Ancestry | SRA Accession |
|---|---|---|---|---|---|
| BZ11-001 | leukocyte | Chunga | 0.0000 | Kinda | SRR7717396 |
| BZ11-002 | leukocyte | Chunga | 0.0000 | Kinda | SRR7717393 |
| BZ11-003 | leukocyte | Chunga | 0.0000 | Kinda | SRR7717394 |
| BZ11-004 | leukocyte | Chunga | 0.0000 | Kinda | SRR7717399 |
| BZ11-005 | leukocyte | Chunga | 0.0000 | Kinda | SRR7717400 |
| BZ11-006 | leukocyte | Chunga | 0.0000 | Kinda | SRR7717397 |
| BZ11-007 | leukocyte | Chunga | 0.0000 | Kinda | SRR7717398 |
| BZ11-008 | leukocyte | Chunga | 0.0000 | Kinda | SRR7717401 |
| BZ11-009 | leukocyte | Chunga | 0.0000 | Kinda | SRR7717402 |
| BZ11-010 | leukocyte | Chunga | 0.0000 | Kinda | SRR7717384 |
| BZ11-011 | leukocyte | Chunga | 0.0000 | Kinda | SRR7717383 |
| BZ11-012 | leukocyte | Chunga | 0.0000 | Kinda | SRR7717386 |
| BZ11-013 | FTA blood spot | Chunga | 0.0000 | Kinda | SRR7717385 |
| BZ11-014 | leukocyte | Chunga | 0.0000 | Kinda | SRR7717388 |
| BZ11-015 | leukocyte | Chunga | 0.0000 | Kinda | SRR7717387 |
| BZ11-016 | leukocyte | Chunga | 0.0000 | Kinda | SRR7717390 |
| BZ11-017 | leukocyte | Chunga | 0.0000 | Kinda | SRR7717389 |
| BZ11-018 | leukocyte | Chunga | 0.0000 | Kinda | SRR7717392 |
| BZ11-019 | leukocyte | Chunga | 0.0000 | Kinda | SRR7717391 |
| BZ11-020 | leukocyte | Chunga | 0.0000 | Kinda | SRR7717293 |
| BZ11-021 | FTA blood spot | Chunga | 0.0000 | Kinda | SRR7717294 |
| BZ11-022 | FTA blood spot | Chunga | 0.0000 | Kinda | SRR7717295 |
| BZ11-023 | FTA blood spot | Chunga | 0.0000 | Kinda | SRR7717296 |
| BZ11-024 | leukocyte | Chunga | 0.0000 | Kinda | SRR7717297 |
| BZ11-025 | leukocyte | Chunga | 0.0000 | Kinda | SRR7717298 |
| BZ11-026 | FTA blood spot | Chunga | 0.0000 | Kinda | SRR7717299 |
| BZ11-028 | leukocyte | Chunga | 0.0000 | Kinda | SRR7717300 |
| BZ11-029 | leukocyte | Chunga | 0.0000 | Kinda | SRR7717301 |
| BZ11-030 | leukocyte | Chunga | 0.0000 | Kinda | SRR7717302 |
| BZ11-031 | leukocyte | Chunga | 0.0000 | Kinda | SRR7717281 |
| BZ11-032 | leukocyte | Chunga | 0.0000 | Kinda | SRR7717280 |
| BZ11-033 | leukocyte | Chunga | 0.0000 | Kinda | SRR7717279 |
| BZ11-034 | leukocyte | Chunga | 0.0000 | Kinda | SRR7717278 |
| BZ11-035 | leukocyte | Chunga | 0.0000 | Kinda | SRR7717277 |
| BZ11-036 | leukocyte | Chunga | 0.0000 | Kinda | SRR7717276 |
| BZ11-037 | leukocyte | Chunga | 0.0000 | Kinda | SRR7717275 |
| BZ11-038 | leukocyte | Chunga | 0.0000 | Kinda | SRR7717274 |
| BZ11-039 | leukocyte | Chunga | 0.0000 | Kinda | SRR7717283 |
| BZ11-040 | leukocyte | Chunga | 0.0000 | Kinda | SRR7717282 |
| BZ11-041 | leukocyte | Chunga | 0.0000 | Kinda | SRR7717321 |
| BZ11-042 | leukocyte | Chunga | 0.0000 | Kinda | SRR7717322 |
| BZ11-043 | FTA blood spot | Chunga | 0.0000 | Kinda | SRR7717319 |
| BZ11-045 | leukocyte | Chunga | 0.0000 | Kinda | SRR7717320 |
| BZ11-046 | FTA blood spot | Chunga | 0.0000 | Kinda | SRR7717317 |
| BZ11-047 | leukocyte | Chunga | 0.0000 | Kinda | SRR7717318 |
| BZ11-048 | FTA blood spot | Chunga | 0.0000 | Kinda | SRR7717315 |
| BZ11-050 | leukocyte | Chunga | 0.0000 | Kinda | SRR7717316 |
| BZ11-051 | FTA blood spot | Chunga | 0.0000 | Kinda | SRR7717313 |
| BZ11-052 | FTA blood spot | Chunga | 0.0000 | Kinda | SRR7717314 |
| BZ11-053 | leukocyte | Chunga | 0.0000 | Kinda | SRR7717310 |
| BZ11-054 | leukocyte | Chunga | 0.0000 | Kinda | SRR7717309 |

continued on next page

Table S1 – continued from previous page

| Sample ID | Sample type | Locality | Hybrid index | Ancestry | SRA Accession |
|---|---|---|---|---|---|
| BZ11-056 | leukocyte | Chunga | 0.0000 | Kinda | SRR7717312 |
| BZ11-057 | leukocyte | Chunga | 0.0508 | hybrid | SRR7717311 |
| BZ11-058 | leukocyte | Chunga | 0.0000 | Kinda | SRR7717306 |
| BZ11-059 | leukocyte | Chunga | 0.0000 | Kinda | SRR7717305 |
| BZ11-061 | FTA blood spot | Chunga | 0.0000 | Kinda | SRR7717308 |
| BZ11-062 | FTA blood spot | Chunga | 0.0000 | Kinda | SRR7717307 |
| BZ11-063 | FTA blood spot | Chunga | 0.0000 | Kinda | SRR7717304 |
| BZ11-064 | FTA blood spot | Chunga | 0.0000 | Kinda | SRR7717303 |
| BZ11-065 | FTA blood spot | Chunga | 0.0000 | Kinda | SRR7717368 |
| BZ11-066 | FTA blood spot | Chunga | 0.0000 | Kinda | SRR7717369 |
| BZ11-067 | FTA blood spot | Chunga | 0.0000 | Kinda | SRR7717370 |
| BZ11-068 | FTA blood spot | Chunga | 0.0000 | Kinda | SRR7717371 |
| BZ11-069 | FTA blood spot | Chunga | 0.0082 | hybrid | SRR7717364 |
| BZ11-070 | FTA blood spot | Chunga | 0.0000 | Kinda | SRR7717365 |
| BZ11-071 | FTA blood spot | Chunga | 0.0000 | Kinda | SRR7717366 |
| BZ11-072 | FTA blood spot | Chunga | 0.0000 | Kinda | SRR7717367 |
| BZ11-073 | FTA blood spot | Chunga | 0.0000 | Kinda | SRR7717361 |
| BZ11-074 | FTA blood spot | Chunga | 0.0000 | Kinda | SRR7717362 |
| BZ11-075 | FTA blood spot | Chunga | 0.0000 | Kinda | SRR7717340 |
| BZ11-076 | FTA blood spot | Chunga | 0.0000 | Kinda | SRR7717334 |
| Chiou-14-030 | feces | Mwengwa Rapids | 0.2039 | hybrid | SRR7717378 |
| Chiou-14-036 | feces | Mwengwa Rapids | 0.2210 | hybrid | SRR7717377 |
| Chiou-14-039 | feces | Mwengwa Rapids | 0.0494 | hybrid | SRR7717380 |
| Chiou-14-041 | feces | Mwengwa Rapids | 0.3463 | hybrid | SRR7717379 |
| Chiou-14-042 | feces | Mwengwa Rapids | 0.0973 | hybrid | SRR7717382 |
| Chiou-14-001 | feces | Malala Camp | 0.5902 | hybrid | SRR7717336 |
| Chiou-14-005 | feces | Malala Camp | 0.4964 | hybrid | SRR7717375 |
| Chiou-14-065 | feces | Malala Camp | 0.3570 | hybrid | SRR7717288 |
| Chiou-14-050 | feces | Musa Bridge | 0.6117 | hybrid | SRR7717373 |
| Chiou-15-003 | feces | Musa Bridge | 0.8600 | hybrid | SRR7717290 |
| Chiou-15-004 | feces | Musa Bridge | 0.4801 | hybrid | SRR7717291 |
| Chiou-15-005 | feces | Musa Bridge | 0.5732 | hybrid | SRR7717292 |
| Chiou-14-054 | feces | Top Musa | 0.8969 | hybrid | SRR7717372 |
| Chiou-14-056 | feces | Top Musa | 0.5515 | hybrid | SRR7717284 |
| Chiou-14-057 | feces | Top Musa | 0.8734 | hybrid | SRR7717285 |
| Chiou-14-058 | feces | Top Musa | 0.7126 | hybrid | SRR7717286 |
| Chiou-14-059 | feces | Top Musa | 0.8396 | hybrid | SRR7717287 |
| BZ07-039 | feces | North Nkala Road | 0.9038 | hybrid | SRR7717331 |
| BZ07-041 | feces | North Nkala Road | 0.6236 | hybrid | SRR7717330 |
| BZ07-042 | feces | North Nkala Road | 1.0000 | chacma | SRR7717329 |
| BZ07-045 | feces | North Nkala Road | 0.8396 | hybrid | SRR7717324 |
| BZ07-047 | feces | North Nkala Road | 0.6173 | hybrid | SRR7717323 |
| Chiou-14-069 | feces | North Nkala Road | 0.6117 | hybrid | SRR7717289 |
| BZ12-001 | plasma | Ngoma Airstrip | 0.5408 | hybrid | SRR7717335 |
| BZ12-002 | plasma | Ngoma Airstrip | 0.7229 | hybrid | SRR7717333 |
| BZ12-003 | plasma | Ngoma Airstrip | 1.0000 | chacma | SRR7717341 |
| BZ12-004 | plasma | Ngoma Airstrip | 0.9415 | hybrid | SRR7717374 |
| BZ12-005 | plasma | Ngoma Airstrip | 0.9498 | hybrid | SRR7717339 |
| BZ12-006 | plasma | Ngoma Airstrip | 1.0000 | chacma | SRR7717338 |
| BZ12-007 | plasma | Ngoma Airstrip | 0.6403 | hybrid | SRR7717363 |
| BZ12-008 | plasma | Ngoma Airstrip | 1.0000 | chacma | SRR7717360 |
| BZ12-009 | plasma | Ngoma Airstrip | 1.0000 | chacma | SRR7717344 |
| BZ12-010 | plasma | Ngoma Airstrip | 0.6125 | hybrid | SRR7717345 |
| BZ12-011 | plasma | Ngoma Airstrip | 0.6172 | hybrid | SRR7717342 |
| BZ12-012 | plasma | Ngoma Airstrip | 0.9511 | hybrid | SRR7717343 |

continued on next page

Table S1 – continued from previous page

| Sample ID | Sample type | Locality | Hybrid index | Ancestry | SRA Accession |
|---|---|---|---|---|---|
| BZ12-030 | plasma | Dendro Park | 1.0000 | chacma | SRR7717348 |
| BZ12-031 | plasma | Dendro Park | 1.0000 | chacma | SRR7717349 |
| BZ12-032 | plasma | Dendro Park | 1.0000 | chacma | SRR7717346 |
| BZ12-033 | plasma | Dendro Park | 1.0000 | chacma | SRR7717347 |
| BZ07-029 | feces | Nanzhila Plains | 1.0000 | chacma | SRR7717328 |
| BZ07-030 | feces | Nanzhila Plains | 0.3894 | hybrid | SRR7717327 |
| BZ07-032 | feces | Nanzhila Plains | 1.0000 | chacma | SRR7717326 |
| BZ07-034 | feces | Nanzhila Plains | 1.0000 | chacma | SRR7717325 |
| BZ07-035 | feces | Nanzhila Plains | 0.3541 | hybrid | SRR7717332 |
| BZ07-004 | feces | Choma | 1.0000 | chacma | SRR7717356 |
| BZ07-005 | feces | Choma | 1.0000 | chacma | SRR7717359 |
| BZ07-007 | feces | Choma | 1.0000 | chacma | SRR7717358 |
| BZ06-218 | feces | Lower Zambezi National Park | 1.0000 | chacma | SRR7717351 |
| BZ06-220 | feces | Lower Zambezi National Park | 1.0000 | chacma | SRR7717350 |
| BZ06-221 | feces | Lower Zambezi National Park | 1.0000 | chacma | SRR7717353 |
| BZ06-225 | feces | Lower Zambezi National Park | 1.0000 | chacma | SRR7717355 |
| BZ06-227 | feces | Lower Zambezi National Park | 1.0000 | chacma | SRR7717354 |

Table S2: Genes with extreme $\alpha_i$.

| Gene ($i$) | Chromosome | Boundaries | | Mean ($\alpha_i$) | ETP interval ($\alpha_i$) | |
|---|---|---|---|---|---|---|
| | | Positive $\alpha_i$ | | | | |
| *RGS12* | 1 | 555231 | 707923 | 1.53853 | 0.12780 | 3.13951 |
| *SLC45A1* | 1 | 11154262 | 11183152 | 1.61807 | 0.41679 | 2.45595 |
| *SLC25A34* | 1 | 18194591 | 18200737 | 1.20260 | 0.10914 | 2.28004 |
| *LDLRAD2* | 1 | 24044439 | 24052967 | 0.90149 | 0.05009 | 1.78411 |
| *LAPTM5* | 1 | 32963659 | 32988931 | 0.77023 | 0.03108 | 1.45115 |
| *C1orf109* | 1 | 39951245 | 39966524 | 1.78544 | 0.60010 | 3.21693 |
| *TESK2* | 1 | 47583596 | 47696639 | 1.41475 | 0.03193 | 2.83091 |
| *PHC3* | 2 | 115535587 | 115621735 | 1.01787 | 0.16558 | 2.19253 |
| *SLC37A3* | 3 | 162125892 | 162193524 | 1.61862 | 0.42885 | 2.89215 |
| *NFKBIE* | 4 | 43632831 | 43641209 | 2.02282 | 0.85927 | 3.49348 |
| uncharacterized | 4 | 74131666 | 74132205 | 0.84749 | 0.04108 | 1.61821 |
| *AKIRIN2* | 4 | 83037303 | 83064962 | 2.08446 | 0.89284 | 3.57656 |
| uncharacterized | 5 | 1127274 | 1400939 | 0.58709 | 0.08970 | 1.05584 |
| *SEMA5A* | 6 | 8812682 | 9007710 | 1.20747 | 0.12999 | 2.19952 |
| *DHX29* | 6 | 51293228 | 51348456 | 1.43984 | 0.24456 | 2.91424 |
| *IK* | 6 | 134169964 | 134181145 | 1.45201 | 0.24013 | 2.71444 |
| *LCTL* | 7 | 40880562 | 40900235 | 1.25034 | 0.26853 | 2.47289 |
| *KIF23* | 7 | 43868409 | 43905208 | 1.99501 | 0.81166 | 3.50669 |
| *FAM149B1* | 9 | 56429241 | 56507955 | 1.38599 | 0.14251 | 2.65677 |
| *PLPP4* | 9 | 112062913 | 112196960 | 1.44887 | 0.25589 | 2.70259 |
| *CFAP46* | 9 | 124314486 | 124435692 | 1.03731 | 0.09627 | 1.88325 |
| *SLC24A3* | 10 | 52238911 | 52441012 | 0.89863 | 0.19719 | 1.68283 |
| *XRN2* | 10 | 54011481 | 54100124 | 1.17762 | 0.21637 | 2.07172 |
| *KIAA1644* | 10 | 84493086 | 84562612 | 1.96598 | 0.52424 | 3.30072 |
| *XPC* | 11 | 82092158 | 82129435 | 1.04051 | 0.05027 | 2.19713 |
| *COQ5* | 11 | 119768501 | 119791403 | 0.95241 | 0.07774 | 1.95525 |
| *TMEM132B* | 11 | 124695755 | 124989612 | 1.40371 | 0.20318 | 2.89844 |
| *STAM2* | 12 | 14362981 | 14415522 | 1.75917 | 0.23931 | 2.84203 |
| *AGAP1* | 12 | 97496718 | 97711301 | 0.59133 | 0.06243 | 1.14762 |
| *UBE2F* | 12 | 99766986 | 99843407 | 2.09450 | 0.80998 | 3.60063 |
| *ALLC* | 13 | 3065128 | 3116554 | 0.80901 | 0.21460 | 1.59779 |
| uncharacterized | 13 | 24314237 | 24346739 | 1.59115 | 0.36145 | 3.07258 |
| *C2orf78* | 13 | 72653999 | 72665448 | 1.22736 | 0.31994 | 2.18866 |
| *TGFBRAP1* | 13 | 96592596 | 96634535 | 1.13548 | 0.28426 | 2.06213 |
| *KDM2A* | 14 | 6836879 | 6978648 | 0.86676 | 0.03077 | 1.82306 |
| *INCENP* | 14 | 11575470 | 11598846 | 1.09960 | 0.03423 | 2.18011 |
| *BCL9L* | 14 | 108226812 | 108247635 | 0.95260 | 0.09708 | 2.21301 |
| *PRDM10* | 14 | 118801938 | 118861158 | 1.32657 | 0.31555 | 2.39229 |
| *NACC2* | 15 | 2212006 | 2260850 | 1.24021 | 0.18861 | 2.16712 |
| *ODF2* | 15 | 9550178 | 9597858 | 1.35574 | 0.20521 | 2.15592 |
| *GOLM1* | 15 | 92537623 | 92608307 | 1.00344 | 0.01040 | 2.23289 |
| *TRPV2* | 16 | 15731378 | 15751538 | 1.75441 | 0.80284 | 2.94720 |
| *PIGL* | 16 | 15853278 | 15962381 | 1.36898 | 0.40052 | 2.56130 |
| *LATS2* | 17 | 140678 | 231436 | 1.73844 | 0.53363 | 2.95804 |
| *RNF138* | 18 | 24060644 | 24097045 | 1.04932 | 0.03257 | 2.33077 |
| *ZNF236* | 18 | 68967234 | 69085587 | 1.05171 | 0.01722 | 2.17271 |
| *TMPRSS9* | 19 | 2175904 | 2187829 | 1.59976 | 0.31975 | 2.50440 |
| *ZNF564* | 19 | 11317666 | 11340206 | 1.01027 | 0.02432 | 2.13787 |
| *ABCC6* | 20 | 15088676 | 15163648 | 0.75447 | 0.08758 | 1.41665 |
| *IQCK* | 20 | 17906074 | 18044544 | 1.44633 | 0.31753 | 2.48762 |
| *PLCG2* | 20 | 63514137 | 63700068 | 1.22891 | 0.39000 | 1.98859 |
| *CBFA2T3* | 20 | 70602695 | 70700524 | 0.65815 | 0.09181 | 1.26826 |

continued on next page

Table S2 – continued from previous page

| Gene ($i$) | Chromosome | Boundaries | | Mean ($\alpha_i$) | ETP interval ($\alpha_i$) | |
|---|---|---|---|---|---|---|
| | | | Negative $\alpha_i$ | | | |
| *AMPH* | 3 | 72517333 | 72771775 | -0.72320 | -1.41438 | -0.05225 |
| *KMT2E* | 3 | 127167177 | 127242094 | -1.11702 | -2.08411 | -0.16206 |
| *LY96* | 8 | 69788797 | 69883096 | -0.47783 | -1.02987 | -0.02982 |

Table S3: Genes with extreme $\beta_i$.

| Gene ($i$) | Chromosome | Boundaries | | Mean ($\beta_i$) | ETP interval ($\beta_i$) | |
|---|---|---|---|---|---|---|
| | | | Positive $\beta_i$ | | | |
| *LIMK1* | 3 | 45157337 | 45194630 | 1.89539 | 0.08140 | 4.03588 |
| *LY96* | 8 | 69788797 | 69883096 | 1.21470 | 0.01039 | 2.65116 |
| *AACS* | 11 | 124402598 | 124484755 | 1.60177 | 0.02148 | 3.38445 |
| *TMEFF2* | 12 | 53718921 | 53960589 | 1.85727 | 0.49071 | 3.47751 |
| *TMEM178A* | 13 | 38564332 | 38615864 | 1.79232 | 0.11404 | 3.94489 |
| | | | Negative $\beta_i$ | | | |
| *ODF2* | 15 | 9550178 | 9597858 | -2.04875 | -4.57410 | -0.01293 |

Table S4: Gene Ontology (GO) terms with significantly enriched $\alpha_i$ or $\beta_i$ cline parameters. Only terms with $p < 0.025$ for any of four one-tailed enrichment tests (positive $\alpha_i$, negative $\alpha_i$, positive $\beta_i$, negative $\beta_i$) are shown here. Key: BP, biological process; CC, cellular component; MF, molecular function; +, test for positive parameter value; –, test for negative parameter value.

| GO term | | Enrichment | | |
|---|---|---|---|---|
| Accession | Name | Parameter | +/– | *p*-value |
| GO:0071902 | positive regulation of protein serine/threonine kinase activity | $\alpha_i$ | + | 0.00123 |
| GO:0046677 | response to antibiotic | $\alpha_i$ | + | 0.00313 |
| GO:0007010 | cytoskeleton organization | $\alpha_i$ | + | 0.00318 |
| GO:0034660 | ncRNA metabolic process | $\alpha_i$ | + | 0.00681 |
| GO:0016192 | vesicle-mediated transport | $\alpha_i$ | + | 0.00718 |
| GO:0060326 | cell chemotaxis | $\alpha_i$ | + | 0.01204 |
| GO:0051258 | protein polymerization | $\alpha_i$ | + | 0.01497 |
| GO:0006508 | proteolysis | $\alpha_i$ | + | 0.01631 |
| GO:0090287 | regulation of cellular response to growth factor stimulus | $\alpha_i$ | + | 0.01711 |
| GO:0033002 | muscle cell proliferation | $\alpha_i$ | + | 0.01896 |
| GO:0006259 | DNA metabolic process | $\alpha_i$ | + | 0.02113 |
| GO:0010595 | positive regulation of endothelial cell migration | $\alpha_i$ | + | 0.02377 |
| GO:0016310 | phosphorylation | $\alpha_i$ | – | 0.00134 |
| GO:0016051 | carbohydrate biosynthetic process | $\alpha_i$ | – | 0.00490 |
| GO:0006650 | glycerophospholipid metabolic process | $\alpha_i$ | – | 0.00579 |
| GO:0008544 | epidermis development | $\alpha_i$ | – | 0.00717 |
| GO:0001501 | skeletal system development | $\alpha_i$ | – | 0.00882 |
| GO:0048706 | embryonic skeletal system development | $\alpha_i$ | – | 0.01223 |
| GO:2001234 | negative regulation of apoptotic signaling pathway | $\alpha_i$ | – | 0.01452 |
| GO:0006355 | regulation of transcription, DNA-templated | $\alpha_i$ | – | 0.01504 |
| GO:0060284 | regulation of cell development | $\alpha_i$ | – | 0.01951 |
| GO:0009306 | protein secretion | $\alpha_i$ | – | 0.01960 |
| GO:0045580 | regulation of T cell differentiation | $\alpha_i$ | – | 0.01967 |
| GO:0002262 | myeloid cell homeostasis | $\alpha_i$ | – | 0.02001 |
| GO:0007009 | plasma membrane organization | $\alpha_i$ | – | 0.02082 |
| GO:0048646 | anatomical structure formation involved in morphogenesis | $\alpha_i$ | – | 0.02142 |
| GO:0030217 | T cell differentiation | $\alpha_i$ | – | 0.02159 |
| GO:0045619 | regulation of lymphocyte differentiation | $\alpha_i$ | – | 0.02364 |
| GO:0030334 | regulation of cell migration | $\beta_i$ | + | 0.00223 |
| GO:0001936 | regulation of endothelial cell proliferation | $\beta_i$ | + | 0.00464 |
| GO:0009967 | positive regulation of signal transduction | $\beta_i$ | + | 0.00527 |
| GO:0007093 | mitotic cell cycle checkpoint | $\beta_i$ | + | 0.00672 |
| GO:0099565 | chemical synaptic transmission, postsynaptic | $\beta_i$ | + | 0.01170 |
| GO:1901991 | negative regulation of mitotic cell cycle phase transition | $\beta_i$ | + | 0.01441 |
| GO:0033674 | positive regulation of kinase activity | $\beta_i$ | + | 0.01452 |
| GO:0045892 | negative regulation of transcription, DNA-templated | $\beta_i$ | + | 0.01556 |
| GO:0007605 | sensory perception of sound | $\beta_i$ | + | 0.01558 |
| GO:0097435 | supramolecular fiber organization | $\beta_i$ | + | 0.01714 |
| GO:0098542 | defense response to other organism | $\beta_i$ | + | 0.02208 |
| GO:0006936 | muscle contraction | $\beta_i$ | + | 0.02307 |
| GO:0016051 | carbohydrate biosynthetic process | $\beta_i$ | – | 0.00497 |
| GO:0003231 | cardiac ventricle development | $\beta_i$ | – | 0.00673 |
| GO:0016570 | histone modification | $\beta_i$ | – | 0.00722 |
| GO:0051209 | release of sequestered calcium ion into cytosol | $\beta_i$ | – | 0.01049 |
| GO:0045732 | positive regulation of protein catabolic process | $\beta_i$ | – | 0.01289 |
| GO:0048525 | negative regulation of viral process | $\beta_i$ | – | 0.01448 |
| GO:0002520 | immune system development | $\beta_i$ | – | 0.01579 |

continued on next page

Table S4 – continued from previous page

| GO term | | Enrichment | | |
|---|---|---|---|---|
| Accession | Name | Parameter | +/– | $p$-value |
| GO:0009887 | animal organ morphogenesis | $\beta_i$ | – | 0.01665 |
| GO:0001568 | blood vessel development | $\beta_i$ | – | 0.01701 |
| GO:0010564 | regulation of cell cycle process | $\beta_i$ | – | 0.02013 |
| GO:0060993 | kidney morphogenesis | $\beta_i$ | – | 0.02138 |
| GO:0006812 | cation transport | $\beta_i$ | – | 0.02172 |
| GO:2000177 | regulation of neural precursor cell proliferation | $\beta_i$ | – | 0.02188 |
| GO:0030001 | metal ion transport | $\beta_i$ | – | 0.02371 |

Table S5: $\beta_i$ parameter values for genes in the JAK/STAT signaling pathway. The point estimate (mean), percentile (relative to point parameter estimates of all genes), and posterior probability of a positive value are shown for $\beta_i$.

| Component | Gene | Chromosome | Point estimate ($\beta_i$) | Percentile | Probability ($\beta_i$ > 0) |
|---|---|---|---|---|---|
| JAK | *JAK1* | 1 | -0.04578 | 41.97 | 0.4744 |
| PIAS | *PIAS1* | 7 | 1.09215 | 98.62 | 0.8550 |
| | *PIAS4* | 19 | 0.86904 | 97.02 | 0.7630 |
| STAT | *STAT2* | 11 | 0.58100 | 92.09 | 0.6786 |
| | *STAT3* | 16 | 0.11558 | 59.80 | 0.5582 |

Table S6: $\alpha_i$ and $\beta_i$ parameter values for genes in the toll-like receptor signaling pathway. The point estimates (means), percentiles (relative to point parameter estimates of all genes), and posterior probabilities are shown for both cline parameters.

| Component | Gene | Chromosome | Point estimate | | Percentile | | Probability | |
|---|---|---|---|---|---|---|---|---|
| | | | $\alpha_i$ | $\beta_i$ | $\alpha_i$ | $\beta_i$ | $\alpha_i > 0$ | $\beta_i < 0$ |
| MKK2 | *MAP2K2* | 19 | 0.64285 | -0.59732 | 92.40 | 7.38 | 0.9450 | 0.8082 |
| IkappaB | *NFKBIE* | 4 | 2.02282 | -1.07868 | 99.91 | 1.91 | 1.0000 | 0.7046 |
